# Sox9^EGFP^ defines biliary epithelial heterogeneity downstream of Yap activity

**DOI:** 10.1101/2020.05.28.113522

**Authors:** Deepthi Y Tulasi, Diego Martinez Castaneda, Kortney Wager, Karel P Alcedo, Jesse R Raab, Adam D Gracz

## Abstract

Intrahepatic bile ducts are lined by biliary epithelial cells (BECs). However, defining the genetic heterogeneity of BECs remains challenging, and tools for identifying BEC subpopulations are limited. Here, we characterize Sox9^EGFP^ transgene expression in the liver and demonstrate that GFP expression levels are associated with distinct cell types. BECs express “low” or “high” levels of GFP, while periportal hepatocytes express “sublow” GFP. Sox9^EGFP^ distribution varies by duct size, with GFP^high^ BECs found at greater numbers in smaller ducts. RNA-seq reveals distinct gene expression signatures for Sox9^EGFP^ populations and enrichment of Notch and Yap signaling in GFP^low^ and GFP^high^ BECs. All GFP^+^ populations are capable of forming organoids, but demonstrate interpopulation differences in organoid survival and size, dependent on media conditions. Organoids derived from Sox9^EGFP^ populations also demonstrate differential activation of HNF4A protein in hepatocyte media conditions, suggesting variable potency in BEC subpopulations. We find that Yap signaling is required to maintain *Sox9* expression in biliary organoids, and that bile acids are insufficient to induce Yap activity or *Sox9 in vivo* and *in vitro*. Our data demonstrate that Sox9^EGFP^ levels provide a readout of Yap activity and delineate BEC heterogeneity, providing a tool for assaying subpopulation-specific cellular function in the liver.

## INTRODUCTION

BECs line intrahepatic bile ducts and are responsible for modifying and transporting bile during homeostasis. Following acute or chronic liver injury, BECs undergo a proliferative response, termed “ductular reaction”, that is associated with liver repair. BECs are also impacted by cholangiopathies, which can result in liver failure and have few therapeutic interventions (Maroni et al. 2015). Direct and indirect evidence suggests a significant degree of functional heterogeneity among BECs with relevance to physiology and liver disease. For example, BEC secretory function is modulated by hormones, peptides, and neurotransmitters, many of which act on a subpopulation of ductal epithelium (Maroni et al. 2015). Recent studies have independently shown that BECs demonstrate differential proliferation following ductal injury, and that some BECs are capable of transdifferentiating into hepatocytes following severe or chronic liver injury (Kamimoto et al. 2016; Raven et al. 2017; Manco et al. 2019). However, assigning potentially heterogeneous responses to specific cell “types” is complicated by the fact that BEC subpopulations lack the clear molecular and genetic definitions that have facilitated a deeper understanding of cell biology in other epithelial tissues. New and accessible tools for interrogating BEC heterogeneity are needed to define and dissect context-dependent roles of BEC subpopulations in liver physiology and disease.

BEC heterogeneity has classically been defined relative to BEC size, with “small” cholangiocytes residing in proximal ductules near the Canals of Herring and “large” cholangiocytes in large ducts (Alpini et al. 1996). While isolated bile ducts and immortalized small and large cholangiocyte cell lines have provided significant insight into biliary physiology, size-based definitions preclude isolation of BECs from genetically- or pharmacologically-challenged livers for direct, *in vivo* insight. Recent single cell transcriptomic studies have shown that BECs demonstrate variable levels of Yes-associated protein (Yap) activity (Pepe-Mooney et al. 2019). Yap activity is increased in bile ducts relative to hepatocytes, and transgenic activation of Yap in hepatocytes drives transdifferentiation to a ductal-phenotype (Yimlamai et al. 2014). As a key regulator of BEC identity, with differential activity across subpopulations of BECs, the Yap pathway and its downstream effectors may present an opportunity for studying BEC heterogeneity from a genetic perspective.

Studies from our lab and others have used a Sox9^EGFP^ BAC transgene to isolate stem, progenitor, and differentiated epithelial cells from mouse intestine and colon (Formeister et al. 2009; Ramalingam et al. 2012; Raab et al. 2020). Because Sox9^EGFP^ is expressed broadly and at distinct levels in these tissues, isolation of cells based on GFP expression provides a single transgene approach to studying cellular heterogeneity. In the liver, *Sox9* is a BEC biomarker that is activated during hepatoblast specification into BEC precursors, where it is required for proper timing of biliary differentiation during development (Antoniou et al. 2009). We hypothesized that Sox9^EGFP^ could facilitate isolation of BEC subpopulations, similar to previous work in the luminal GI tract. Here, we examine Sox9^EGFP^ transgene expression in intrahepatic bile ducts, and exploit differential GFP expression levels to isolate distinct cellular subpopulations. Our results demonstrate that Sox9^EGFP^ expression levels facilitate dissection of BEC heterogeneity.

## RESULTS

### Sox9^EGFP^ is expressed in intrahepatic BECs and periportal hepatocytes

We sought to determine if the Sox9^EGFP^ BAC transgene, previously established as a stem/progenitor cell marker in intestinal and colonic epithelium, accurately labels known *Sox9*^+^ populations in the liver. Low magnification, epifluorescent imaging of whole liver lobes demonstrated robust GFP signal in branching patterns consistent with intrahepatic bile ducts (Fig. 1A). Examination of histological sections revealed GFP^+^ cells with typical ductal morphology confined to the portal area and co-localized with endogenous SOX9 (Supplemental Fig. 1A). We also observed Sox9^EGFP^ expression throughout the biliary tree, including the gallbladder, extrahepatic bile duct, and pancreatic duct, consistent with known expression patterns of *Sox9* in these tissues (Supplemental Fig. 1B-F). Due to independent developmental origins and significant differences in gene expression of biliary epithelium in extrahepatic tissue, we focused the present study on intrahepatic ducts (Zong and Stanger 2012; Rimland et al. 2020).

**Figure 1.**
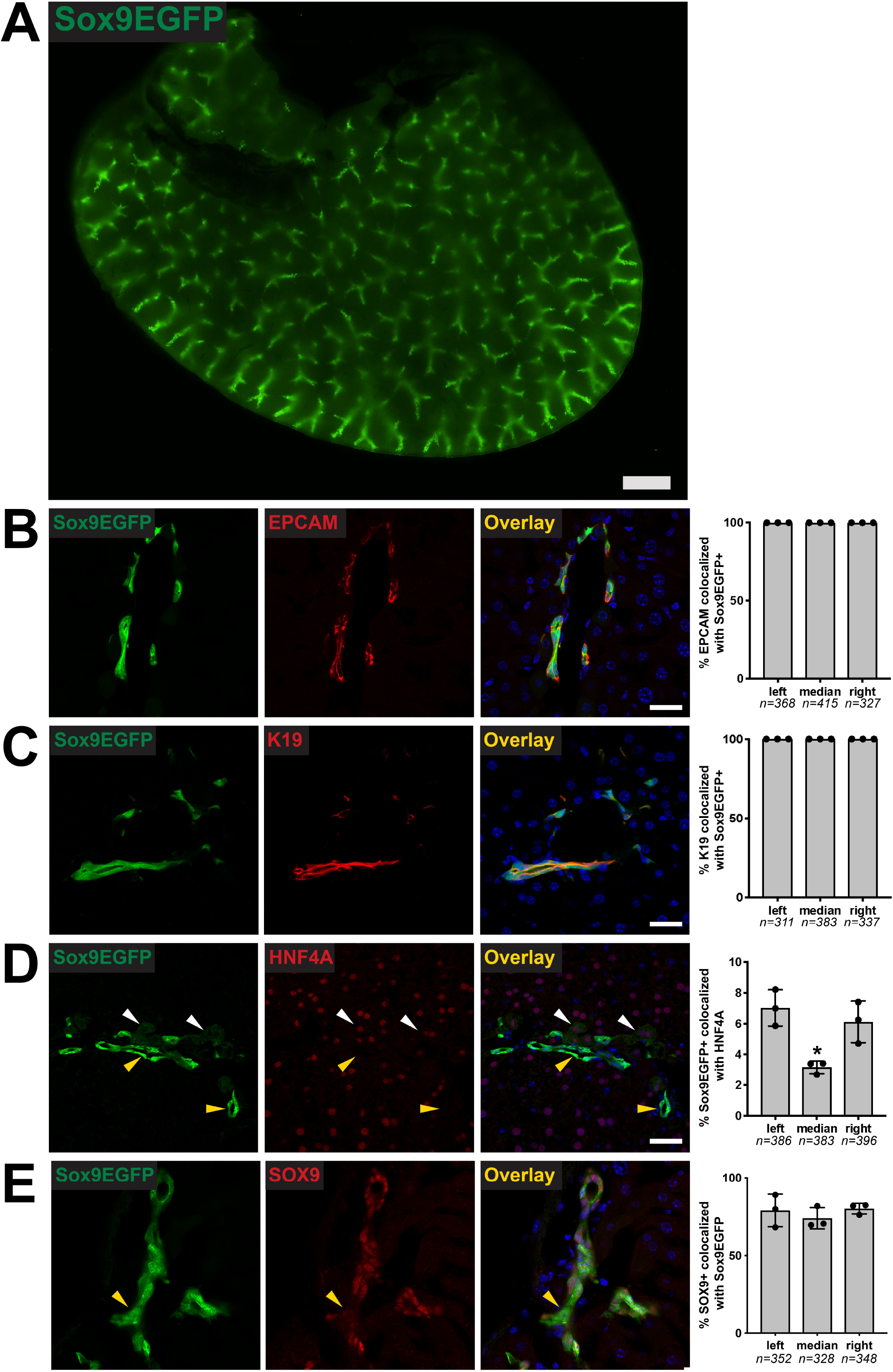
Sox9^EGFP^ is expressed in intrahepatic bile ducts and peribiliary hepatocytes. (A) Low-magnification imaging of Sox9^EGFP^ expression throughout the intrahepatic biliary tree (right lobe, scale bar = 1mM). Immunofluorescence demonstrates that EGFP is co-expressed in (B) 100% of EPCAM^+^ and (C) K19^+^ positive BECs across left, median, and right lobes. (D) Rare peribiliary cells expressing very low levels of EGFP co-localize with HNF4A and are morphologically consistent with hepatocytes (white arrows indicate EGFP^+^/HNF4A^+^, yellow arrows indicate EGFP^+^/HNF4A^−^). (E) SOX9 is expressed in most, but not all, Sox9^EGFP+^ cells (yellow arrows indicate EGFP^+^/SOX9^−^) (scale bars = 50μm; * indicates p < 0.05, one-way ANOVA and Tukey’s test).

To determine: (1) whether Sox9^EGFP^ expression accurately labels all BECs and (2) if Sox9^EGFP^ is restricted to BECs, we next examined EGFP expression relative to the independent BEC markers EPCAM and cytokeratin 19 (K19) by immunofluorescence. In both cases, we found that 100% of BECs identified by EPCAM or K19 were also positive for GFP in left, median, and right lobes (Fig. 1B, C). Previous reports have shown that a subpopulation of periportal hepatocytes co-express BEC markers, including *Sox9* (Font-Burgada et al. 2015). To test if Sox9^EGFP^ is also expressed in hybrid hepatocytes, we co-localized GFP+ cells with hepatocyte transcription factor HNF4A. We found that a small percentage of Sox9^EGFP^ cells are HNF4A+ (Fig. 1D). These cells were observed to have typical hepatocyte morphology and express very low levels of the GFP transgene (Fig. 1D, white arrowheads), in contrast to GFP^+^ cells with BEC morphology, which were appreciably brighter (Fig. 1D, yellow arrowheads). We did not observe any co-localization between Sox9^EGFP^ and vimentin, which is expressed on mesenchymal, endothelial, and stellate cells (Supplemental Fig. 1G) (Troeger et al. 2012).

Finally, we co-localized Sox9^EGFP^ with endogenous SOX9. Surprisingly, we found that some cells expressing the GFP transgene did not co-express endogenous SOX9 protein (Fig. 1E, arrows). The number of GFP+/SOX9-cells was not significantly variable between biological replicates (Supplemental Fig. 1H). Because SOX9 is considered a pan-biliary marker, we independently confirmed the presence of SOX9-BECs by co-localizing SOX9 with EPCAM by immunofluorescence. We found EPCAM+/SOX9-BECs (79.0% ± 10.5%), at approximately the same rate as EGFP+/SOX9-BECs (77.7% ± 7.1%) (Supplemental Fig. 2B). Collectively, our immunofluorescence analyses confirm that Sox9^EGFP^ is expressed ubiquitously in BECs, including BECs that lack endogenous SOX9 protein. Additionally, very low levels of GFP are expressed in a subpopulation of periportal hepatocytes, consistent with known expression of endogenous *Sox9* in “hybrid” hepatocytes.

**Figure 2.**
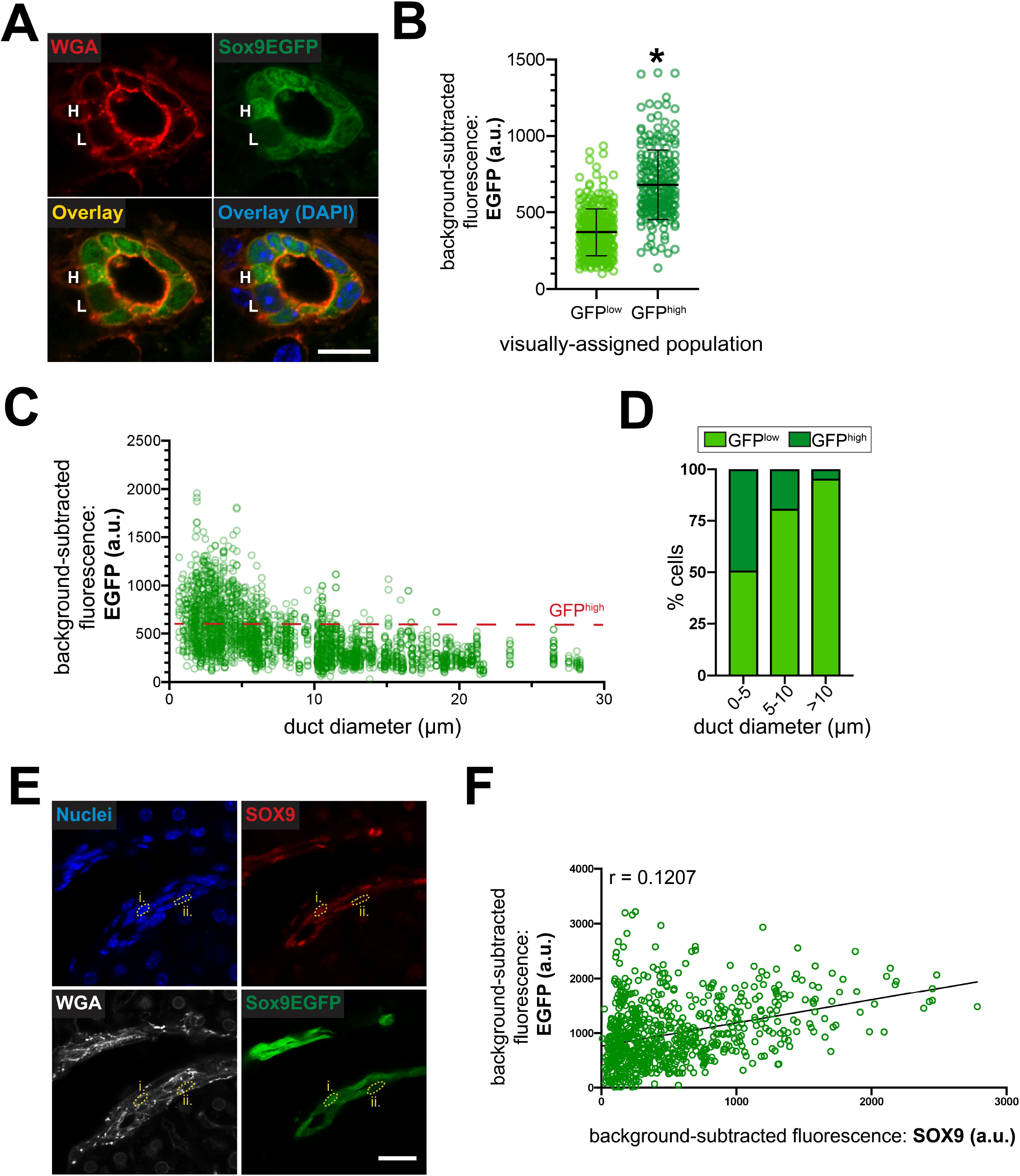
GFP^high^ BECs are enriched in small intrahepatic bile ducts. (A) WGA labels cell membranes and facilitates quantification of Sox9^EGFP^ cells qualitatively identified as “high” (H) and “low” (L) (scale bar = 10μm). (B) Qualitatively-identified GFP^high^ cells are significantly brighter than GFP^low^ cells by quantification of confocal images (n=307 GFP^low^, 215 GFP^high^; * indicates p < 0.001, unpaired t-test; a.u. = arbitrary units) (C) Quantification of EGFP intensity relative to duct diameter demonstrates the distribution of Sox9^EGFP^ populations across the intrahepatic biliary tree (n=2,589 cells) (D) GFP^high^ BECs are most abundant in small ducts and rare with increasing duct size (n=954 cells 0-5μm, 523 cells 5-10μm, 1,112 cells >10μm). (E) Bisbenzimide and WGA label nuclei and cell membranes respectively for quantification of SOX9 and EGFP in single cells (roman numerals denote cells across multiple channels; scale bar = 25μm). (F) SOX9 correlates poorly with EGFP levels (n=747 cells).

### GFP^high^ cells are more plentiful in smaller ducts

While qualitative observation demonstrated variable Sox9^EGFP^ expression in intrahepatic bile ducts, we sought to quantify ductal GFP at the single cell level and determine if different levels of expression correlate with anatomical localization. We employed semi-quantitative confocal microscopy and measured GFP in individual cells using wheat germ agglutin (WGA) to delineate cell membranes. To avoid artifacts associated with antibody detection, all experiments measured endogenous GFP. First, we visually categorized BECs as GFP^low^ or GFP^high^ and asked if qualitatively identified Sox9^EGFP^ populations demonstrated quantitatively discernible differences in GFP intensity (Fig. 2A). We found that BECs identified as GFP^high^ had significantly higher fluorescence intensity relative to those identified as GFP^low^, validating our ability to resolve Sox9^EGFP^ populations by histology (Fig. 2B).

Because BEC heterogeneity has been classically described relative to cell location in small or large ducts, we next quantified GFP expression in individual BECs relative to duct diameter (Alpini et al. 1996). GFP fluorescence and duct diameter of the resident bile duct were measured for 2,589 BECs, and we noted a clear inverse relationship between the number of GFP^high^ cells and duct diameter (Fig. 2C). To quantify the percentage of GFP^high^ BECs in different sized ducts, we arbitrarily defined GFP^high^ BECs as having a mean fluorescence intensity ≥600, based on the upper limit of fluorescence in a majority of GFP^low^ cells (Fig. 2B). We determined that the smallest bile ducts (0-5μm luminal diameter) are comprised of 49.16% GFP^high^ and 50.84% GFP^low^ cells, intermediate ducts (5-10μm luminal diameter) are 18.93% GFP^high^ and 81.07% GFP^low^ cells, and the largest ducts (>10μm luminal diameter) are comprised of 4.50% GFP^high^ and 95.50% GFP^low^ cells (Fig. 2D).

Sox9^EGFP^ is driven by upstream transcriptional regulators of *Sox9*, and GFP levels accurately predict endogenous SOX9 protein levels in GFP+ cells of the small intestine and colon (Formeister et al. 2009; Ramalingam et al. 2012). We quantified 747 BECs, using WGA and bisbenzimide as membrane and nuclear markers for GFP and SOX9 immunofluorescence, respectively (Fig. 2E). GFP level was a poor predictor of endogenous SOX9 protein level and we did not observe specific association of SOX9-cells with low or high GFP expression (Fig. 2F). Interestingly, some BECs expressing the highest observed level of GFP expressed little or no appreciable SOX9. Together, these data demonstrate that Sox9^EGFP^ BECs are compartmentalized by duct size, with GFP^high^ cells present in increased numbers in smaller ducts. Further, we find that Sox9^EGFP^ level is not predictive of SOX9 protein expression, in contrast with observations in the small intestine and colon.

### Sox9^EGFP^ expression levels facilitate isolation of cellular subpopulations

To further quantify relative levels of Sox9^EGFP^ expression, we analyzed intrahepatic GFP by flow cytometry. Livers were dissociated using a protocol optimized to obtain single BECs, at the expense of hepatocyte viability (Broutier et al. 2016). Flow cytometry revealed that a majority of cells in our single cell isolations were negative for the GFP transgene (88.7% ± 6.1%) (Fig. 3A, Supplemental Fig. 2A&B). Within the GFP+ fraction, we defined three distinct subpopulations: GFP^sublow^ (GFP^sub^), GFP^low^, and GFP^high^ (Fig. 3A&B). Of GFP^+^ populations, GFP^high^ cells (71.0% ± 6.1%) were significantly more abundant than GFP^low^ (11.9% ± 1.8%) and GFP^sub^ (10.6% ± 4.4%) cells (Supplemental Fig. 2C).

**Figure 3.**
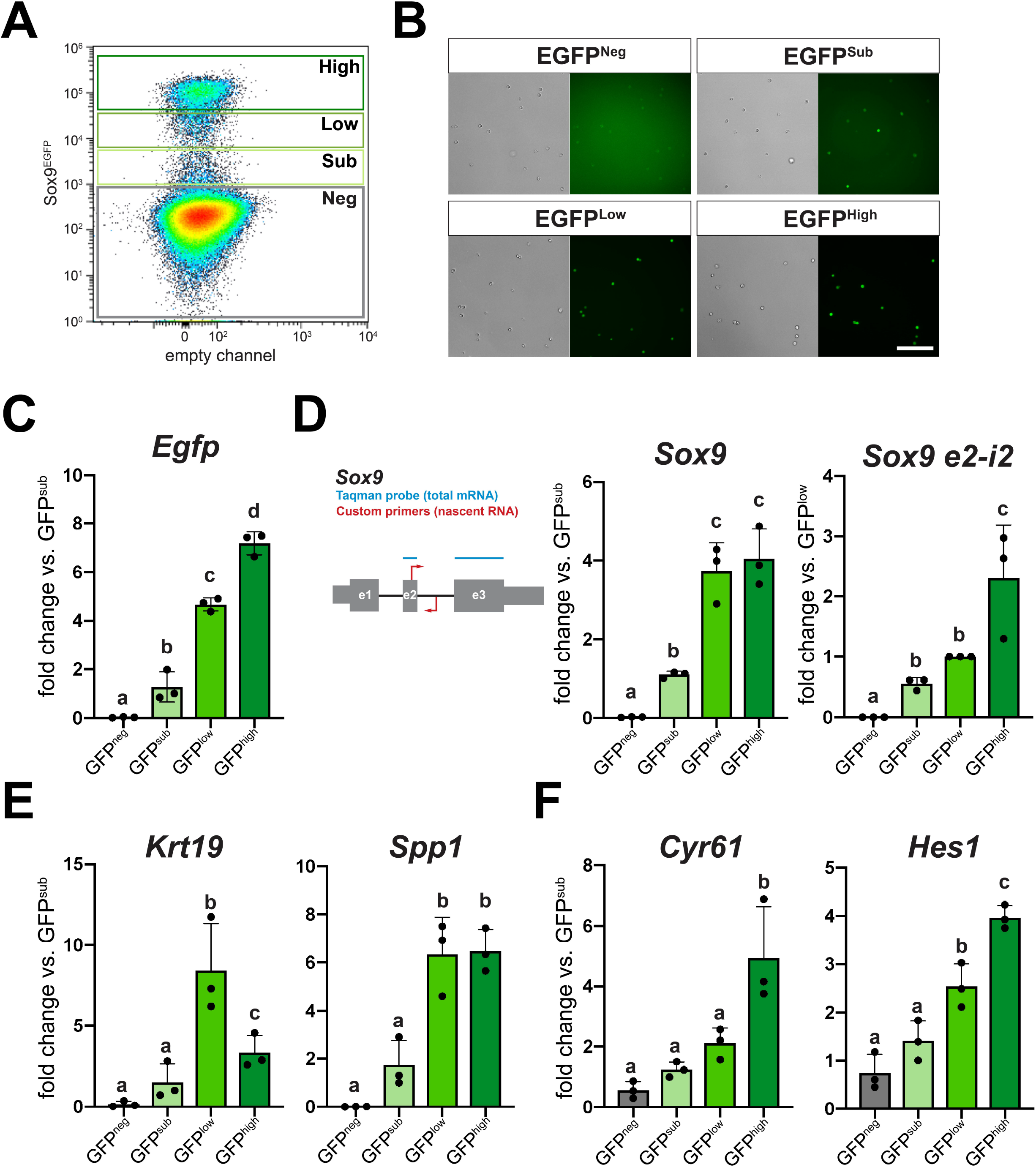
FACS isolation identifies Sox9^EGFP^ populations with distinct gene expression patterns. (A) Sox9^EGFP^ is divided into four populations by flow analysis. (B) FACS-isolated cells from Sox9^EGFP^ populations exhibit increasing EGFP intensity (scale bar = 100μm). (C) *Egfp* expression is significantly different between GFP^neg, sub, low,^ and ^high^ populations. (D) RT-qPCR probes detecting *Sox9* mRNA show enrichment between GFP^neg^ and ^sub^ populations, but not GFP^low^ and ^high^. Primers against nascent *Sox9* RNA demonstrate upregulation in GFP^high^. (E) Canonical BEC biomarkers *Krt19* and *Spp1* are differentially express in Sox9^EGFP^ populations. (F) *Cyr61* and *Hes1* are most significantly enriched in GFP^high^ cells, suggesting increased Yap activity (letters indicate grouping by significance, p < 0.05, one-way ANOVA and Tukey’s test).

Based on our histological assays, we reasoned that GFP^low^ and GFP^high^ populations were most likely to represent cells of the intrahepatic bile ducts, while GFP^sub^ were likely to represent periportal hepatocytes. Because biliary heterogeneity has classically been defined relative to BEC size, we next asked if cells isolated from GFP^low^ and GFP^high^ populations differed in area (Alpini et al. 1996). We found that, on average, GFP^high^ cells were significantly smaller than GFP^low^ cells, consistent with previous models of small and large cholangiocytes (Supplemental Fig. 2D). These data demonstrate that intrahepatic Sox9^EGFP^ is expressed at distinct levels that can be resolved by flow cytometry, and suggest that these levels are associated with BEC subpopulations previously defined by cell size.

### Sox9^EGFP^ populations exhibit differential gene expression patterns

We next analyzed gene expression in FACS-isolated Sox9^EGFP^ BEC populations. As expected, *Egfp* was differentially expressed across GFP^neg^, GFP^sub^, GFP^low^, and GFP^high^ populations (Fig. 3C). *Sox9* was enriched as expected between: (1) GFP^neg^ and GFP^sub^ and (2) GFP^sub^ and GFP^low/high^ (Fig. 3D). However, we observed no difference in *Sox9* expression between GFP^low^ and GFP^high^ populations (Fig. 2D). We reasoned that differences in post-transcriptional regulation could lead to differential *Egfp* expression without differential *Sox9* expression. To test this hypothesis, we designed RT-qPCR primers spanning the second exon and intron of Sox9 in order to detect nascent *Sox9* RNA and complement our Taqman assay, which spanned exons 2 and 3 of *Sox9* and is specific to mRNA (Fig. 3D). We found that GFP^high^ cells express significantly higher levels of nascent *Sox9* RNA relative to GFP^low^ and GFP^sub^ populations (Fig. 3D). These data demonstrate that Sox9^EGFP^ levels provide a readout of *Sox9* transcription, which is highest in GFP^high^ BECs despite equivalent amounts of *Sox9* mRNA between GFP^low^ and GFP^high^ populations. This also suggests that *Sox9* undergoes post-transcriptional regulation in BEC subpopulations.

Next, we examined the expression of canonical BEC genes in Sox9^EGFP^ populations. *Spp1*, which encodes pan-biliary marker osteopontin (OPN), was significantly enriched in both GFP^low^ and GFP^high^ populations (Fig. 3E). Interestingly, while both GFP^low^ and GFP^high^ cells expressed significantly higher levels of *Krt19* relative to GFP^neg^ and GFP^sub^ populations, we observed enrichment of *Krt19* in GFP^low^ relative to GFP^high^ (Fig. 3E). To test if Sox9^EGFP^ populations capture recently-reported Yap-associated heterogeneity, we assayed expression of *Cyr61* and *Hes1*, which have been previously identified as YAP target genes and proposed biomarkers of BEC heterogeneity (Yimlamai et al. 2014; Pepe-Mooney et al. 2019). Both genes were significantly upregulated in GFP^high^ cells relative to GFP^low^ cells, and *Hes1* was also enriched in GFP^low^ relative to GFP^sub^ (Fig. 3F). These data support histological evidence that GFP^low^ and GFP^high^ cells represent BEC populations, imply distinct transcriptional identities for GFP^low^ and GFP^high^ BECs, and suggest that Sox9^EGFP^ expression levels capture previously described heterogeneity relative to ductal YAP activity.

### Unique transcriptomic signatures define intrahepatic Sox9^EGFP^ populations

To determine gene expression signatures of Sox9^EGFP^ subpopulations, we conducted RNA-seq on FACS-isolated GFP^neg^, GFP^sub^, GFP^low^, and GFP^high^ populations. Clustering by principal components analysis (PCA) reinforced expected relationships predicted by histology and RT-qPCR. GFP^low^ and GFP^high^ populations, which are both consistent with BEC identity, clustered together while GFP^neg^ and GFP^sub^ samples demonstrated more significant differences (Fig. 4A). Differential gene expression analysis identified genes unique to and shared between Sox9^EGFP^ populations, with the largest shared gene sets consisting of genes shared between GFP^neg^ and GFP^sub^ populations, followed by genes shared between GFP^low^ and GFP^high^ populations (Fig. 4B). By comparison, very few genes were shared between GFP^low/high^ and GFP^sub^, and no genes were shared between GFP^low/high^ and GFP^neg^.

**Figure 4.**
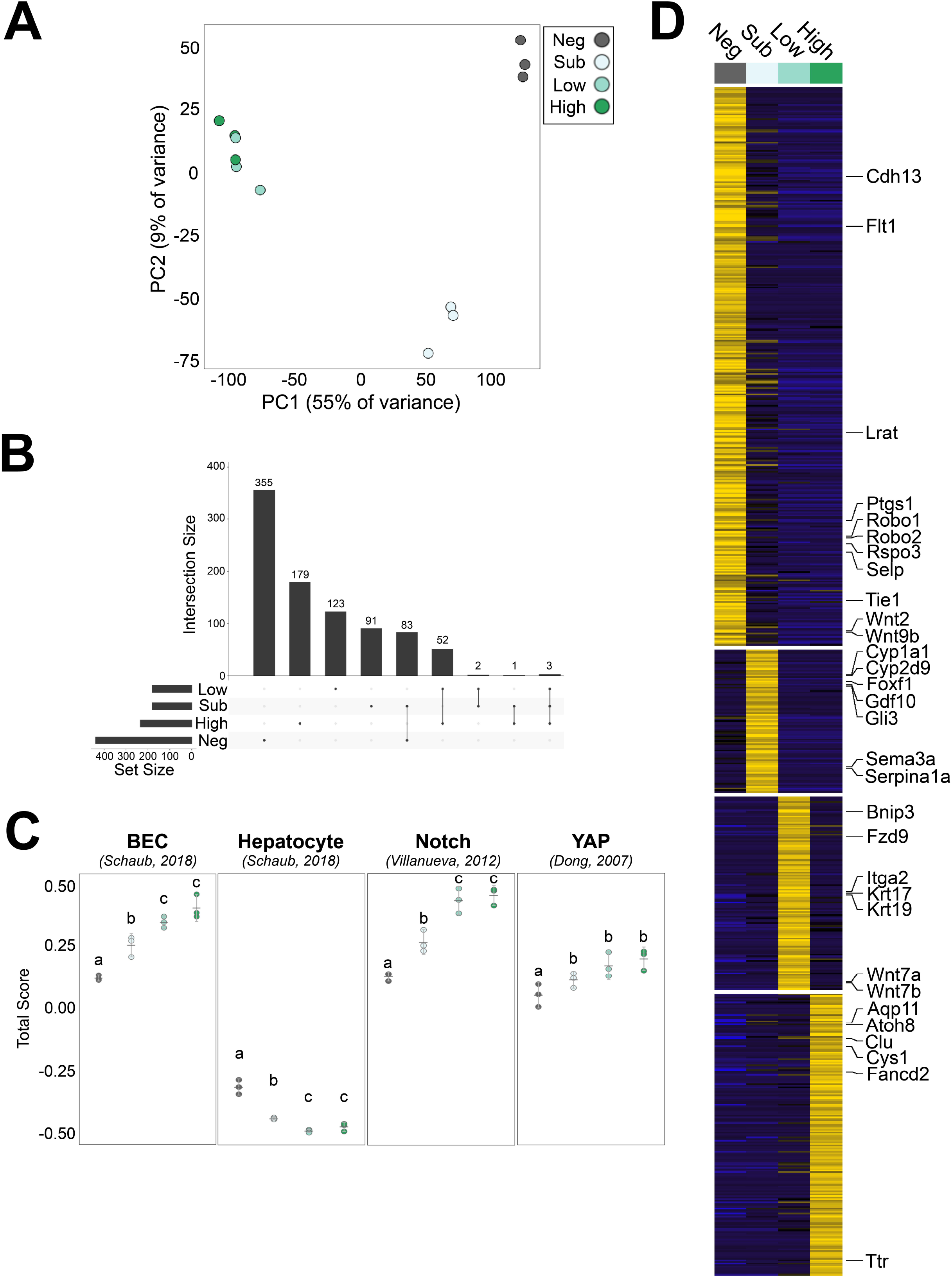
Sox9^EGFP^ populations have distinct transcriptomes. (A) GFP^neg^ and ^sub^ populations form distinct clusters by PCA, while GFP^low^ and ^high^ cluster similarly. (B) Genes shared between multiple populations reinforce similarities between GFP^low^ and ^high^. (C) Gene signature analysis demonstrates significant enrichment of BEC, Notch, and Yap gene sets in GFP^low^ and ^high^ populations relative to GFP^neg^ and ^sub^, while all populations are depleted for hepatocyte genes (error bars represent 95% CI; letters indicate grouping by significance, p < 0.05, one-way ANOVA and Tukey’s test). (D) Heatmap represents genes uniquely upregulated in a single Sox9^EGFP^ population.

We next analyzed gene expression signatures of Sox9^EGFP^ populations against published transcriptomic datasets using rank-based gene set scoring (Foroutan et al. 2018). Gene signature analysis demonstrated that GFP^low^ and GFP^high^ were more enriched for intrahepatic BEC-associated genes than GFP^neg^ and GFP^sub^ (Fig. 4C) (Schaub et al. 2018). All four populations were depleted for genes associated with hepatocytes, affirming that our single cell isolation and collection protocol selects against GFP^neg^ hepatocytes. To examine enrichment of known biliary regulatory pathways, we assayed Notch and Yap target genes and again observed significant enrichment in GFP^low^ and GFP^high^ populations (Fig. 4C) (Dong et al. 2007; Villanueva et al. 2012). While our qPCR data demonstrated differential expression of specific YAP target genes, transcriptome-scale analyses did not reveal a significant difference between GFP^low^ and GFP^high^ populations for either Yap or Notch gene signature scores (Fig. 4C).

To more stringently assay transcriptomic differences between Sox9^EGFP^ populations, we identified genes that were significantly upregulated in a single population (Fig. 4D and Supplemental Table 1). Many of the genes uniquely enriched in the GFP^neg^ population were consistent with pericentral liver sinusoidal endothelial cells (LSECs), including: *Rspo3, Cdh13*, *Flt1*, *Ptgs1*, *Selp, Wnt2*, and *Wnt9b* (Fig. 4D) (Halpern et al. 2018). While pericentral LSECs express CD31, it remains restricted to the cytoplasm, explaining why this population would persist in our isolation strategy despite negative selection against CD31 by FACS (DeLeve et al. 2004) (Supplemental Fig. 2A). The hepatic stellate cell-specific gene, *Lrat*, was also upregulated in GFP^neg^ cells (Mederacke et al. 2013). While a majority of genes specific to GFP^+^ populations have no established function in the liver, we observed gene expression patterns consistent with hybrid hepatocyte and BEC identities. Hepatocyte-associated metabolic enzymes *Cyp1a1* and *Serpina1a* were both upregulated in GFP^sub^ cells, along with the Hedgehog signaling target *Gli3*. Consistent with our qPCR data, *Krt19* was significantly enriched in GFP^low^ BECs, along with *Krt17* and WNT pathway genes *Fzd9*, *Wnt7a*, and *Wnt7b*. GFP^high^ BECs expressed *Aqp11*, previously shown to be enriched in developing bile ducts, and *Clu*, which was recently identified as a marker and functional mediator of facultative stem cells in the small intestine (Fig. 4D) (Poling et al. 2014; Ayyaz et al. 2019). Together, our transcriptomic data reinforce the broad cellular identities of Sox9^EGFP^ populations defined by histology and RT-qPCR, and identify unique gene expression signatures that differentiate GFP^sub^ hybrid hepatocytes and GFP^low^ and GFP^high^ BECs.

### Sox9^EGFP^ populations exhibit distinct phenotypic responses to single-cell organoid culture

To assay functional properties of proliferation and cell fate in Sox9^EGFP^ populations, we turned to organoid assays. Liver organoids were initially reported to be formed by BECs, with subsequent studies describing media conditions supportive of hepatocyte organoids (Huch et al. 2013; Peng et al. 2018). Since the Sox9^EGFP^ transgene is expressed in both a subpopulation of hepatocytes (GFP^sub^) as well as BECs (GFP^low^ and GFP^high^), we used conditions developed for BECs (BEC media) as well as conditions developed for hepatocyte organoid culture (TNFα media). All GFP^+^ populations formed organoids in both media conditions, while GFP^neg^ cells never formed organoids (Fig. 5A). Consistent with previous reports, organoids with clear lumens or spherical morphology took approximately 5-6 days to form, and we quantified organoid formation rates at 7 and 14 days of culture (Aloia et al. 2019). TNFα media increased organoid formation in all GFP^+^ populations, relative to BEC media (Supplemental Fig. 5A). While all GFP^+^ populations formed organoids at approximately the same rate in BEC media, single GFP^low^ and GFP^high^ cells grown in TNFα media formed significantly more organoids than GFP^sub^ cells at both 7 and 14 days of culture (Fig. 5B). We observed an increase in GFP^low^-derived organoids between days 7 and 14, suggesting an extended delay in the ability of some GFP^low^ BECs to form morphologically appreciable organoids in TNFα media (Fig. 5B).

**Figure 5.**
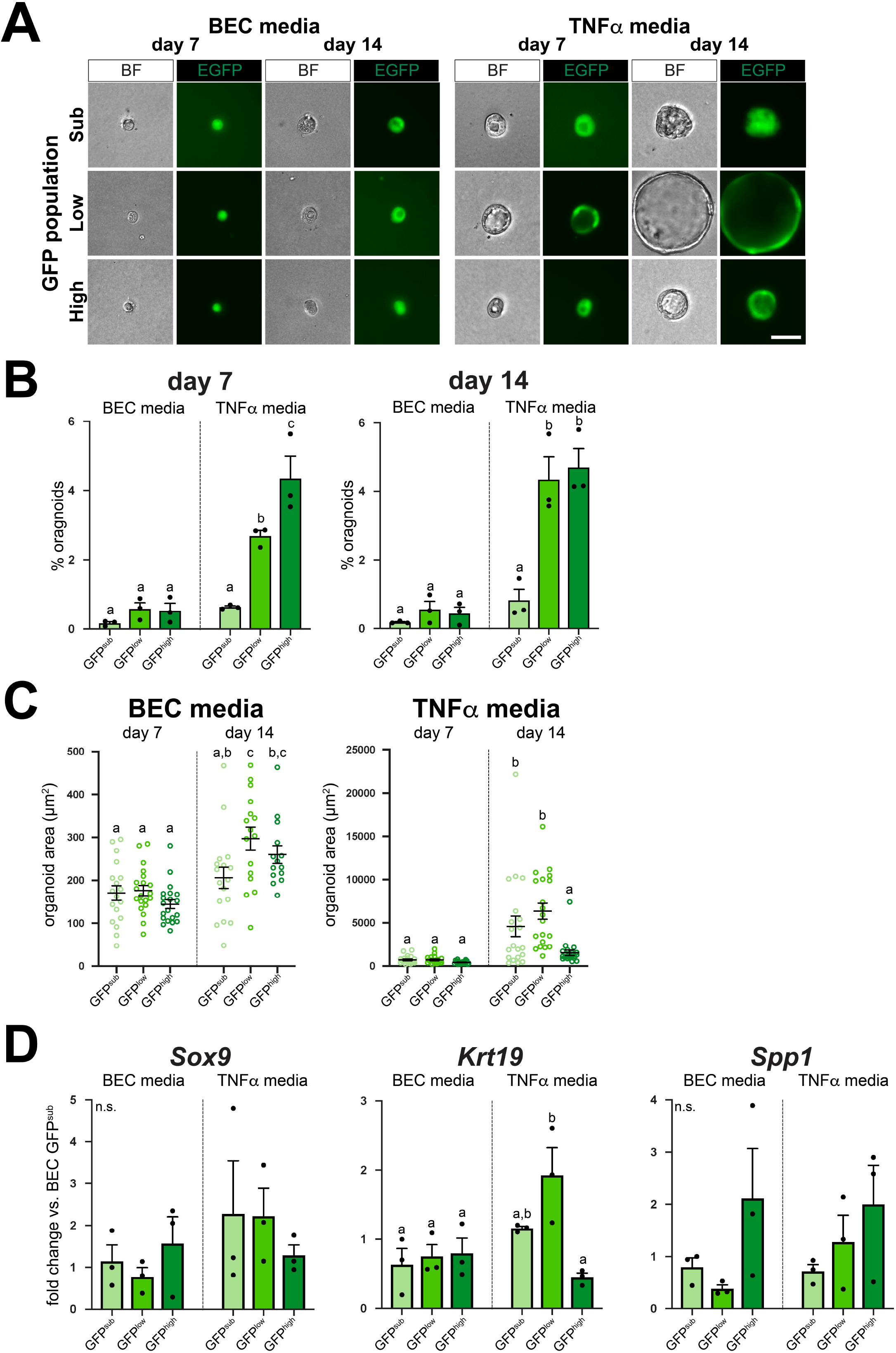
Functional differences in organoids derived from single GFP^+^ cells are dependent on media conditions. (A) Single cells from Sox9^EGFP^ populations form organoids in media conditions developed for BECs (BEC media) and hepatocytes (TNFα media) (scale bar = 25 μm). (B) TNFα media significantly increases organoid formation relative to BEC media in all Sox9^EGFP^ populations, and are required for significantly different organoid formation rates between GFP^sub, low,^ and ^high^. (C) TNFα media also drives more significant increases in organoid size between 7 and 14 days of culture and results in significantly larger GFP^sub^ and ^low^-derived organoids at day 14 relative to GFP^high^-derived organoids at the same timepoint. (D) *Sox9* and *Spp1* are not differentially expressed in BEC or TNFα media, but *Krt19* is enriched in GFP^low^-derived organoids relative to GFP^high^-derived organoids in TNFα media only, exhibiting an expression pattern similar to *in vivo* results (letters indicate grouping by significance, p < 0.05, one-way ANOVA and Tukey’s test).

Next, we quantified organoid area and found that organoids grown in TNFα media were significantly larger at both timepoints and across Sox9^EGFP^ populations (Supplemental Fig. 5B). Further, we found that relative to organoids grown in BEC media, organoids grown in TNFα media demonstrated a greater increase in average area between days 7 and 14 in culture, suggesting more robust proliferation induced by TNFα media (Fig. 5C). Additionally, GFP^sub^- and GFP^low^-derived organoids were significantly larger than GFP^high^-derived organoids at day 14 in TNFα media (Fig. 5C).

To determine if organoids derived from different Sox9^EGFP^ populations exhibited distinct gene expression profiles, we conducted qPCR on organoids grown in BEC and TNFα media for 7 days. While we observed some trends in expression for most genes examined, variability between biological replicates resulted in few significant differences between Sox9^EGFP^ populations or media conditions. While *Sox9* and *Spp1* were not differentially expressed, *Krt19* remained upregulated in GFP^low^-derived organoids grown in TNFα media relative to all organoids grown in BEC media and GFP^high^-derived organoids grown in TNFα media (Fig. 5D). *Cyr61* was enriched in GFP^high^-derived BEC media organoids relative to GFP^sub^-derived TNFα media, but not significantly different between other populations and conditions (Supplemental Fig. 5C). Additionally, *Klf6* trended toward enrichment in GFP^low^-derived organoids grown in both media conditions (Supplemental Fig. 5C). Despite increased organoid size and the use of WNT agonist CHIR99021 in TNFα media, Wnt targets *Ccnd1* and *Myc* were not significantly upregulated in organoids grown in TNFα media (Supplemental Fig. 5D). Our results demonstrate that organoid-forming capacity is restricted to Sox9^EGFP^ populations, but suggest that GFP^sub^, GFP^low^, and GFP^high^ cells perform similarly in organoid assays in terms of survival and gene expression. Our data also demonstrate that the most pronounced differences in organoid survival and size were found between media conditions rather than Sox9^EGFP^ populations. Further, significant inter-population differences between organoid survival, size, and *Krt19* expression were found exclusively in TNFα media, suggesting that functional heterogeneity of BECs may be driven by exogenous conditions.

### Sox9^EGFP^ populations demonstrate different rates of transdifferentiation

Previous reports have indicated that BECs are capable of transdifferentiating into hepatocytes, and vice-versa, in defined organoid culture systems (Huch et al. 2013; Hu et al. 2018). Since TNFα media conditions were developed for long-term hepatocyte organoid culture, we reasoned that BEC vs. hepatocyte fate decisions might differ between BEC media and TNFα media. To assay cellular identity, we examined protein expression of SOX9 and hepatocyte-specific transcription factor HNF4A by whole-mount immunofluorescence in single cell-derived Sox9^EGFP^ organoids following 7 days of culture in BEC or TNFα media (Fig. 6A). Hepatocytes were isolated by collagenase perfusion and grown in TNFα media as positive controls for hepatocyte identity and HNF4A expression (Supplemental Fig. 4A). Organoids produced by single GFP^sub^ and GFP^low^ cells expressed SOX9 exclusively in BEC media, while a small number of GFP^high^-derived organoids (8.0%) co-expressed SOX9 and HNF4A (Fig. 6B and Supplemental Fig. 4C). Interestingly, rare GFP^high^-derived organoids were also observed to be negative for both SOX9 and HNF4A, consistent with SOX9^−^/GFP^+^ BECs observed *in vivo*.

**Figure 6.**
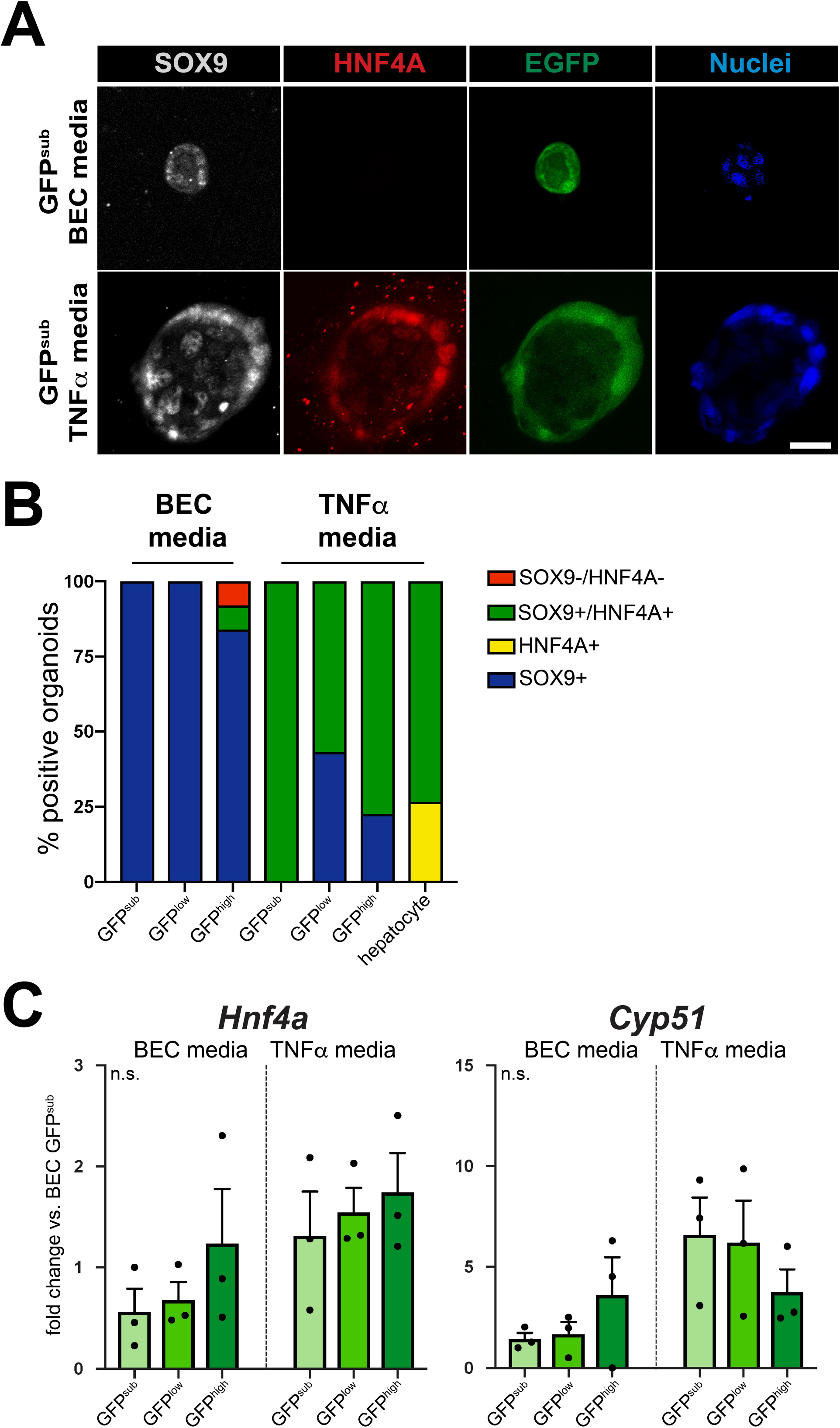
Differential rates of HNF4A activation in Sox9^EGFP^ populations. (A) Representative images of SOX9 and HNF4A immunofluorescence in organoids derived from single Sox9^EGFP^ populations (scale bar = 25 μm). (B) SOX9 and HNF4A expression is dependent on both media conditions and Sox9^EGFP^ population, with TNFα media driving increased expression of HNF4A in GFP^sub, low,^ and ^high^-derived organoids, but at distinct rates (n per group in Supplemental Fig. 4C). (C) Despite increased and differential HNF4A activation across Sox9^EGFP^ populations in TNFα media, *Hnf4a* and *Cyp51* gene expression is not changed.

Single cells grown in TNFα media produced organoids that were more likely to express HNF4A relative to single cells grown in BEC media, regardless of which Sox9^EGFP^ population they were derived from (Fig. 6B). Strikingly, while GFP^sub^-derived organoids did not express HNF4A in BEC media, 100% co-expressed SOX9 and HNF4A in TNFα media. SOX9 and HNF4A were also co-expressed in a larger proportion of GFP^high^-derived (77.4%) organoids in TNFα media versus GFP^low^ (56.8%) (Fig. 6B). Only collagenase-isolated hepatocytes produced organoids expressing HNF4A in the absence of SOX9. Next, we examined gene expression of *Hnf4a* and *Cyp51*, which was previously reported to be upregulated in Lgr5^EGFP+^ liver organoids in hepatocyte media conditions (Huch et al. 2013). In contrast to HNF4A protein expression, neither gene was differentially expressed across Sox9^EGFP^ populations or media conditions. These data demonstrate that organoids derived from Sox9^EGFP^ populations exhibit varying potential to express HNF4A and that HNF4A expression is enhanced in all Sox9^EGFP^ populations by TNFα media conditions. Our gene expression data suggests that HNF4A activation in TNFα media is regulated at the post-transcriptional level.

### Sox9 expression is maintained by Yap, but is not modulated by bile acids

*Sox9* is known to be activated downstream of Yap activity in diverse cellular contexts, including developing and regenerating hepatocytes and esophageal cancer cells (Song et al. 2014; Yi et al. 2016). Since we observed upregulation of *Cyr61* and *Hes1* in GFP^high^ cells, we hypothesized that Sox9^EGFP^ may serve as a readout for Yap in BECs. To test if biliary *Sox9* expression is Yap-dependent, we treated organoids isolated from whole bile ducts with verteporfin, a small molecule inhibitor of Yap (Liu-Chittenden et al. 2012). Verteporfin treatment did not result in appreciable changes to organoid morphology (Fig. 7A). As expected, we observed a significant decrease in the expression of Yap target genes *Cyr61* and *Klf6*, as well as *Sox9* (Fig. 7B). *Gadd45b*, another known target of Yap signaling, remained unchanged, suggesting differential regulation of canonical Yap targets in biliary epithelium (Fig. 7B).

**Figure 7.**
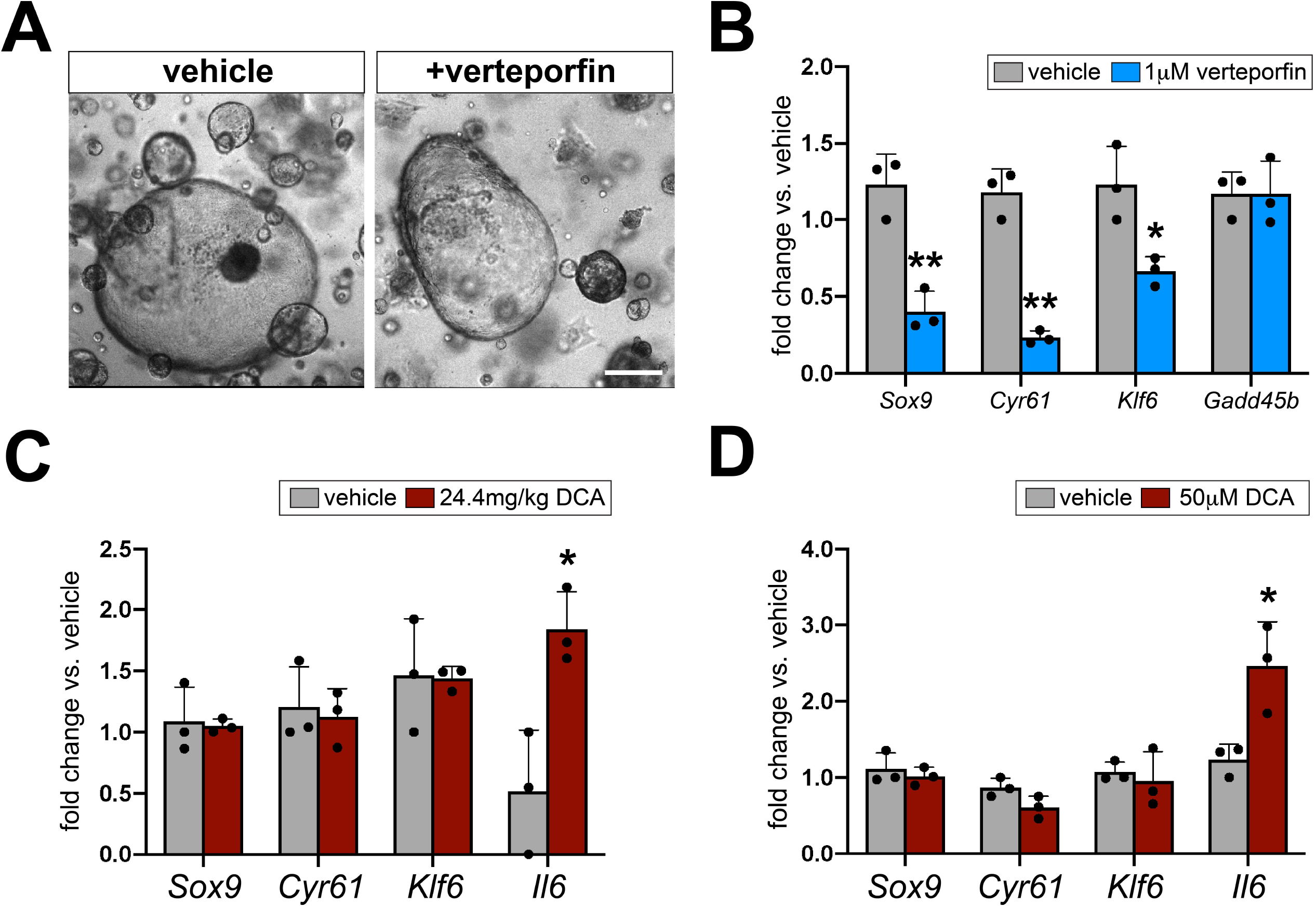
Sox9 expression is maintained by Yap and independent of bile acid signaling. (A) Verteporfin treatment does not impact organoid morphology (scale bar = 50μm), but (B) results in significant downregulation of *Sox9* and established BEC Yap targets, *Cyr61* and *Klf6*, while *Gadd45b* is unaffected. (C) *Sox9* and Yap target genes are not differentially expressed in EPCAM^+^ cells from mice treated with DCA for 24hr, but *Il6* is significantly upregulated. (D) Primary biliary monolayers treated with DCA *in vitro* reproduce *in vivo* results, upregulating *Il6*, but failing to demonstrate significantly different expression of *Sox9*, *Cyr61*, or *Klf6* (* indicates significance, p < 0.05; ** indicates significance, p ≤ 0.01, unpaired t-test).

Recent studies have shown that bile acids induce Yap in both hepatocytes and BECs (Anakk et al. 2013; Pepe-Mooney et al. 2019). To test if biliary *Sox9* is also induced by bile acids upstream of Yap, we administered deoxycholic acid (DCA) to wild-type mice via a single, intraperitoneal injection and analyzed gene expression in FACS-isolated, EPCAM^+^ BECs 24hr later (Paolini et al. 2002) (Supplemental Fig. 7A). We found that neither *Sox9* or Yap targets, *Cyr61* or *Klf6*, were responsive to DCA *in vivo* (Fig. 7C). To identify a BEC-specific positive control gene and validate DCA treatment, we assayed a panel of putative targets of farnesoid X receptor (FXR) that were: (1) identified by published FXR ChIP-seq, and (2) expressed in GFP^low^ or GFP^high^ BECs in our RNA-seq (Thomas et al. 2010). While most candidate target genes were unresponsive to DCA, we observed significant upregulation of *Il6* in DCA-treated mice relative to vehicle controls, consistent with previous reports of NF-λB induction of *Il6* in BECs following bile acid treatment (Dai et al. 2013).

We reasoned that transcriptional responses to bile acid fluctuations *in vivo* might be subtle, and next assayed BEC-autonomous responses to DCA by treating organoids *in vitro*. We did not observe a change in expression of *Sox9* or Yap target genes at 24hr of treatment, despite induction of *Il6* in 2 of 3 replicates (Supplemental Fig. 5C). Because DCA-induced Yap was previously shown to be dependent on ASBT, which is found on the apical membrane of BECs, we confirmed that DCA was interacting with apical cell surfaces via two independent experiments (Pepe-Mooney et al. 2019). First, we treated biliary organoids with FITC-conjugated DCA and measured luminal fluorescence 9hr after treatment. Organoids treated with FITC-DCA exhibited significantly increased luminal fluorescence relative to controls treated with FITC-dextran, indicating that DCA is capable of crossing the basolateral membrane and entering organoid lumens (Supplemental Fig. 7B). Next, we passaged biliary organoids into monolayers, so apical membranes interact directly with overlaid media. As observed in our *in vivo* and organoid experiments, DCA failed to induce *Cyr61*, *Klf6*, or *Sox9* in biliary monolayers, but *Il6* was significantly upregulated (Fig. 7D). These data demonstrate that biliary *Sox9* expression is maintained by Yap, but that both Yap activity and *Sox9* are unaffected by DCA.

## DISCUSSION

Dissecting cellular heterogeneity is important for understanding how subpopulations of cells contribute to tissue homeostasis and disease. While advances in single cell genomics provide tools for rapid and unbiased cataloguing of cell types, models that facilitate identification and manipulation of cell populations in intact tissues are still critical for understanding cellular function. Here, we characterize Sox9^EGFP^ expression in the liver, and establish a tool for understanding heterogeneity in BECs. Sox9^EGFP^ is expressed at variable levels that are associated with phenotypically distinct cell populations in the small intestine, colon, and pancreatic duct (Formeister et al. 2009; Ramalingam et al. 2012; Rezanejad et al. 2018). Our data demonstrate variability in Sox9^EGFP^ expression levels throughout the intrahepatic bile ducts and periportal hepatocytes, and confirm unique anatomical distribution, gene expression, and functional properties characteristic of each Sox9^EGFP^ population. We also observe distinct Sox9^EGFP^ expression levels in the extrahepatic bile ducts and gallbladder, suggesting that the transgene may be a useful biomarker for understanding cellular heterogeneity in these tissues as well.

Unlike previous reports in the intestine, we find that Sox9^EGFP^ does not accurately report endogenous SOX9 protein or mRNA levels. Instead, the highest levels of *Egfp* expression are associated with increased nascent *Sox9* RNA. Additionally, some GFP^+^ BECs do not express SOX9 protein. Since SOX9 is considered a ubiquitous marker of BECs and is required for BEC specification during development, this observation was unexpected, but confirmed by co-localization of EPCAM and SOX9 demonstrating a similar number of SOX9^−^ BECs (Antoniou et al. 2009). The mechanistic significance of both the regulation of *Sox9* mRNA and protein expression, as well as its impact on adult BEC identity and function warrants further investigation.

In contrast to other epithelial tissues, the biliary epithelium lacks clearly defined cell populations with compartmentalized functional properties (for e.g.: stem, enteroendocrine, and absorptive cells in intestinal epithelium; type I and type II pneumocytes in alveolar epithelium). This lack of defined populations and general ambiguity of distinctions between cell types in the biliary epithelium complicates defining cell types captured within each Sox9^EGFP^ population, as well as intrapopulation heterogeneity. The clearest distinction found in the present study is between GFP^sub^ hepatocytes and GFP^low/high^ BECs. *Sox9* expressing “hybrid” hepatocytes have been previously defined by their peribiliary anatomical location and co-expression of hepatocyte and BEC markers (Font-Burgada et al. 2015). GFP^sub^ cells are localized to the peribiliary niche, co-express HNF4A *in vivo*, and demonstrate enrichment of BEC genes by transcriptomic signature analysis, consistent with hybrid hepatocytes. Interestingly, GFP^sub^-derived organoids exclusively expressed SOX9 in BEC media and always co-expressed SOX9 and HNF4A in TNFα media. While the low rate of organoid formation precluded functional assays in the present study, these data suggest bipotency of GFP^sub^ cells and their progeny.

Transcriptomic differences are present between GFP^low^ and GFP^high^ BEC populations, but are more subtle. Our data demonstrate that GFP^high^ BECs resemble previously defined small cholangiocytes, based on their anatomical enrichment in small, proximal ducts, as well as their size relative to GFP^low^ BECs (Alpini et al. 1996). Interestingly, we find that GFP^low^ BECs express significantly higher levels of *Krt19*, a pan-biliary marker that has been associated with BEC maturation in induced pluripotent stem cell models (Sampaziotis et al. 2015). However, it is unclear if differential *Krt19* expression *in vivo* is indicative of different stages of hierarchical cell fate, or simply heterogeneity across distinct, mature cell types. Our RNA-seq analysis also uncovered *Wnt7a* and *Wnt7b*, which were identified as specific to a subpopulation of BECs in a DDC damage model, as uniquely enriched in GFP^low^ cells (Pepe-Mooney et al. 2019). Though we did not detect a significant difference in a previously published Yap gene signature between GFP^low^ and GFP^high^ populations, we observed increased expression of *Cyr61* and *Hes1* in GFP^high^ vs. GFP^low^ BECs by RT-qPCR (Dong et al. 2007). *Cyr61* and *Hes1* have previously been identified as Yap-regulated markers of BEC heterogeneity (Pepe-Mooney et al. 2019). Taken in context of our verteporfin experiments demonstrating dependence of *Sox9* expression on Yap signaling in organoids, these results suggest that high levels of Sox9^EGFP^ expression in BECs are associated with increased Yap activity.

In organoid-forming assays using media conditions developed for biliary organoids, both GFP^low^ and GFP^high^ BECs behaved similarly in terms of organoid size and survival (Huch et al. 2013). It is known that despite heterogeneous Yap activity *in vivo*, BEC media conditions drive broad activation of Yap *in vitro*, which could lead to homogenization of organoid phenotypes (Pepe-Mooney et al. 2019). Compellingly, we find that TNFα media conditions result in organoid size differences between GFP^low^- and GFP^high^-derived organoids, as well as increased initial organoid formation from GFP^high^ cells. This may indicate that exogenous factors drive or repress functional heterogeneity in BECs. Like GFP^sub^ cells, GFP^low^ and GFP^high^ cells also demonstrated differential activation of HNF4A in TNFα media. Interestingly, GFP^high^-derived organoids were more likely to express HNF4A, tempting speculation that different subpopulations of BECs may be more likely to undergo transdifferentiation to hepatocytes. Because organoid culture conditions impact multiple signaling pathways and fail to perfectly recapitulate the *in vivo* environment, further work will be required to expand insight into how *in vitro* conditions affect BEC subpopulations.

Future studies may benefit from scRNA-seq to further tease apart heterogeneity within Sox9^EGFP^ populations. However, the lack of distinct clustering of total BEC scRNA-seq by t-Distributed Stochastic Neighbor Embedding (t-SNE) suggests that deep sequencing would be required to detect differences between subpopulations driven by low-expressed genes (Pepe-Mooney et al. 2019; Planas-Paz et al. 2019). Our organoid transdifferentiation studies, which demonstrate changes in HNF4A protein expression despite no significant change in *Hnf4a* gene expression, along with the presence of SOX9-BECs, suggest that post-transcriptional regulation may be important in biliary identity. Proteomic characterization may also be required to further resolve BEC heterogeneity. Additionally, we show that inter-population differences in organoid size and survival are found in TNFα media conditions but not in BEC media. This suggests that BEC functional heterogeneity, which may be associated with emergent transcriptomic and proteomic heterogeneity, could be context-dependent and revealed by changes in the larger biliary “niche”. The Sox9^EGFP^ model presented here will provide a platform for further exploration of such heterogeneity in biliary homeostasis, injury, and regeneration.

---

**Supplemental Figure 1.**
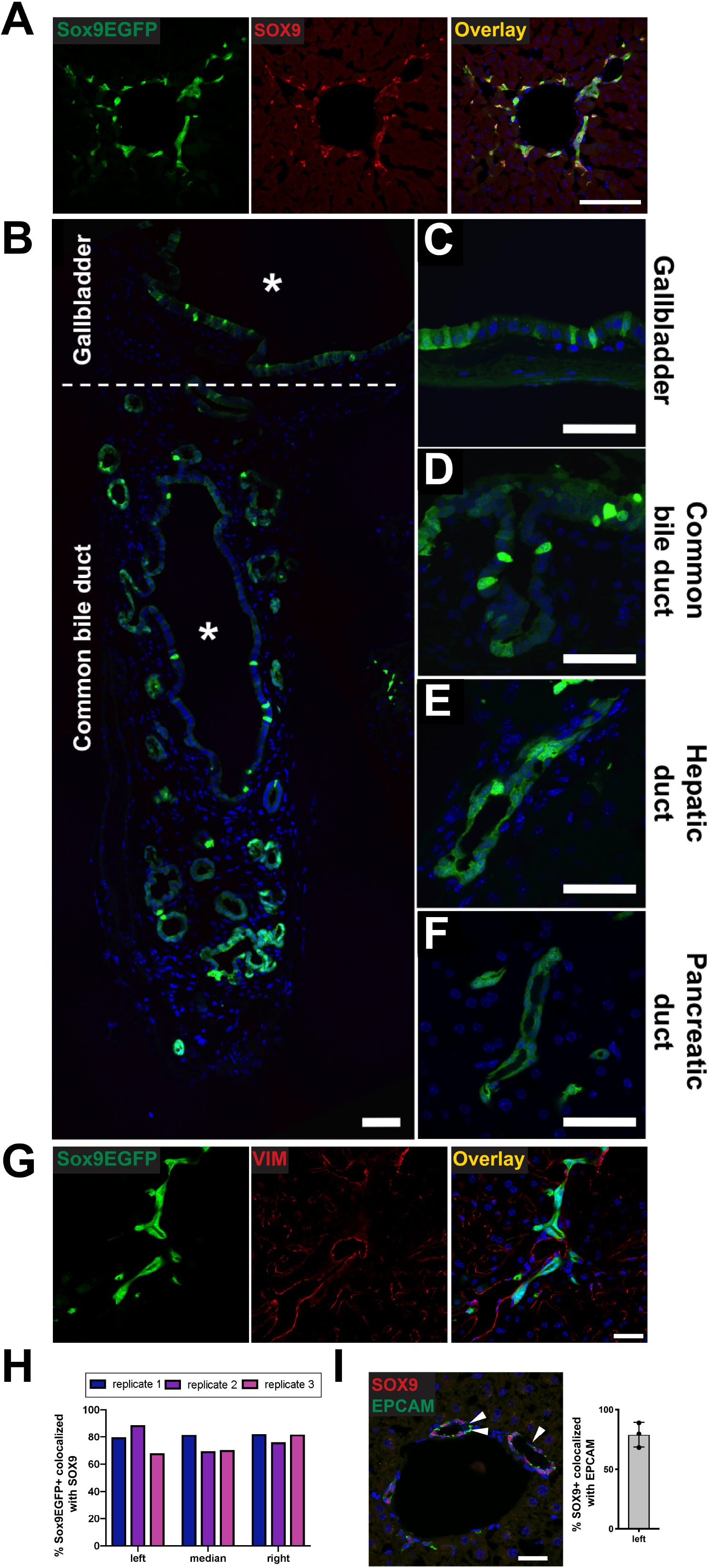
Sox9^EGFP^ transgene expression is variable throughout the biliary tree. (A) Sox9^EGFP^ is expressed in intrahepatic bile ducts and co-localizes with SOX9 (scale bar = μm). (B) Epithelial cells of the (C) gallbladder, (D) common bile duct, (E) hepatic ducts, and (F) pancreatic ducts express Sox9^EGFP^ at variable levels (asterisks indicate lumens; scale bars = 5μm). (G) Sox9^EGFP^ does not co-localize with mesenchymal marker, vimentin (VIM) (scale bar = 5μm). (H) SOX9^−^/EGFP^+^ BECs are found consistently across liver lobes and biological replicates. (I) Co-localization of SOX9 with EPCAM independently confirms the presence and occurrence of SOX9-BECs (left lobe, arrows indicate EPCAM+/SOX9-cells, scale bar = 5μm).

**Supplemental Figure 2.**
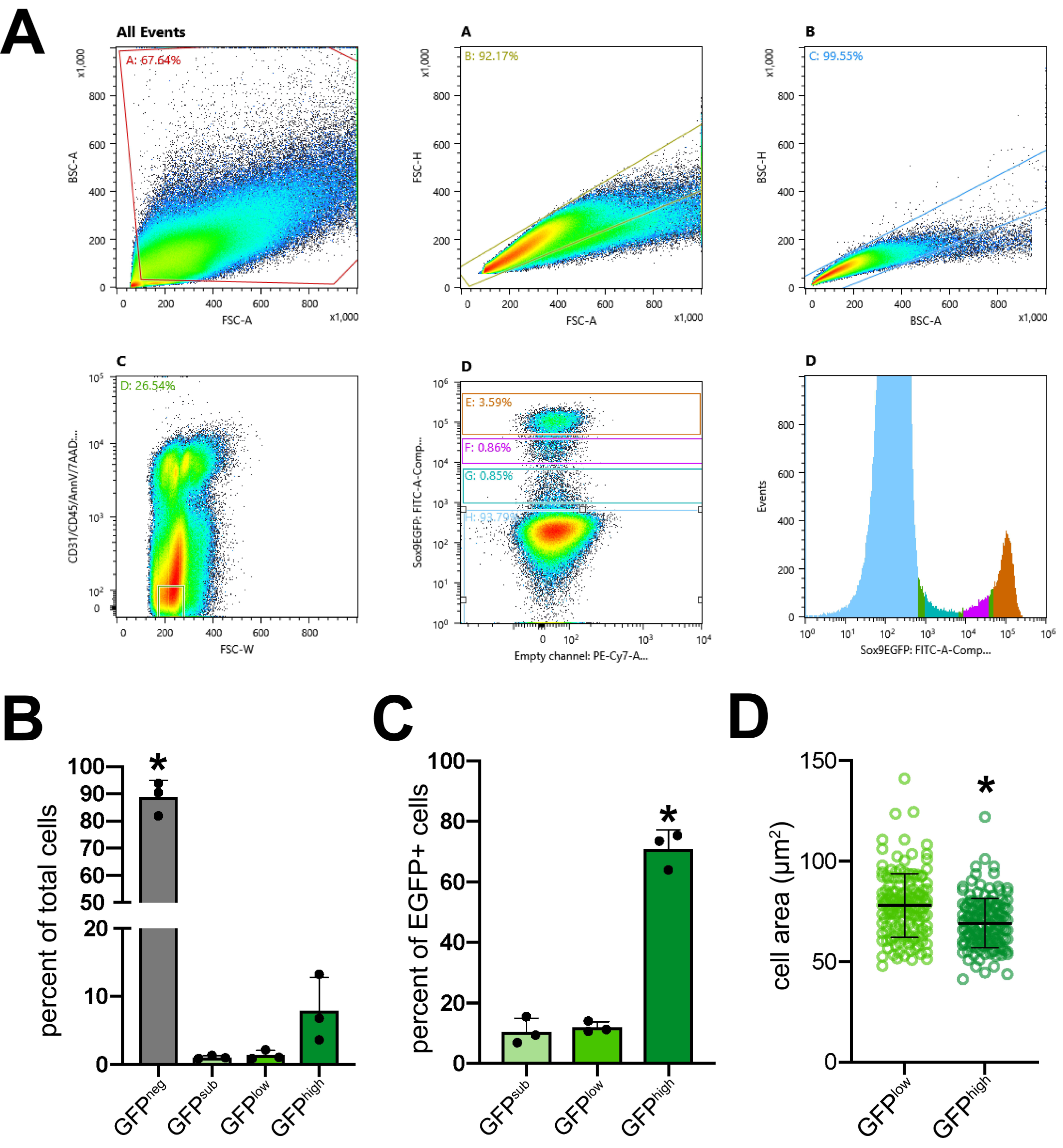
FACS isolation of Sox9^EGFP^ populations. (A) FACS isolation strategy for Sox9^EGFP^ populations. (B) Flow analysis demonstrates that GFP^neg^ cells are the most abundant population present in single cell preps after exclusion of CD31/CD45/Annexin V/7-AAD^+^ cells, and (C) GFP^high^ cells are the most abundant of GFP^+^ populations (* indicates significance, p < 0.05, one-way ANOVA and Tukey’s test). (D) Cell area measurements demonstrate that isolated GFP^high^ cells are significantly smaller than GFP^low^ cells (n=150 GFP^low^, 150 GFP^high^; * indicates p < 0.001, unpaired t-test).

**Supplemental Figure 3.**
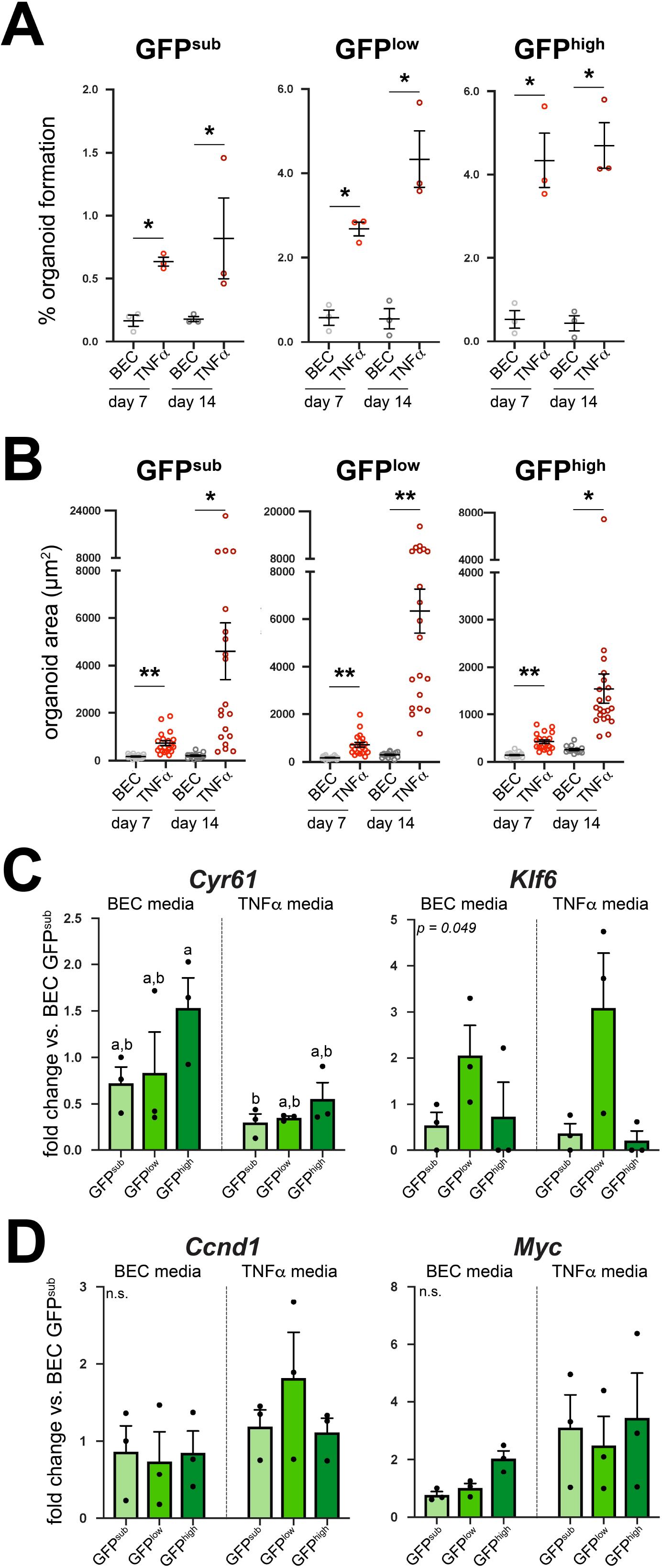
Survival and size of organoids derived from single Sox9^EGFP+^ cells. TNFα media results in increased organoid formation rates relative to BEC media at 7 and 14 days of culture in all Sox9^EGFP^ populations (* indicates significance, p < 0.01, unpaired t-test). TNFα media results in increased organoid size relative to BEC media at 7 and 14 days of culture in all Sox9^EGFP^ populations (* indicates significance, p < 0.002; ** indicates significance, p ≤ 0.0001, unpaired t-test). (C) *Cyr61* is significantly upregulated in GFP^high^-derived BEC media organoids relative to GFP^sub^-derived TNFα media. While *Klf6* is not significantly regulated between populations and conditions, it trends toward increased expression in GFP^low^-derived organoids in both BEC and TNFα media. (D) WNT target genes *Myc* and *Ccnd1* are not differentially expressed between populations and conditions.

**Supplemental Figure 4.**
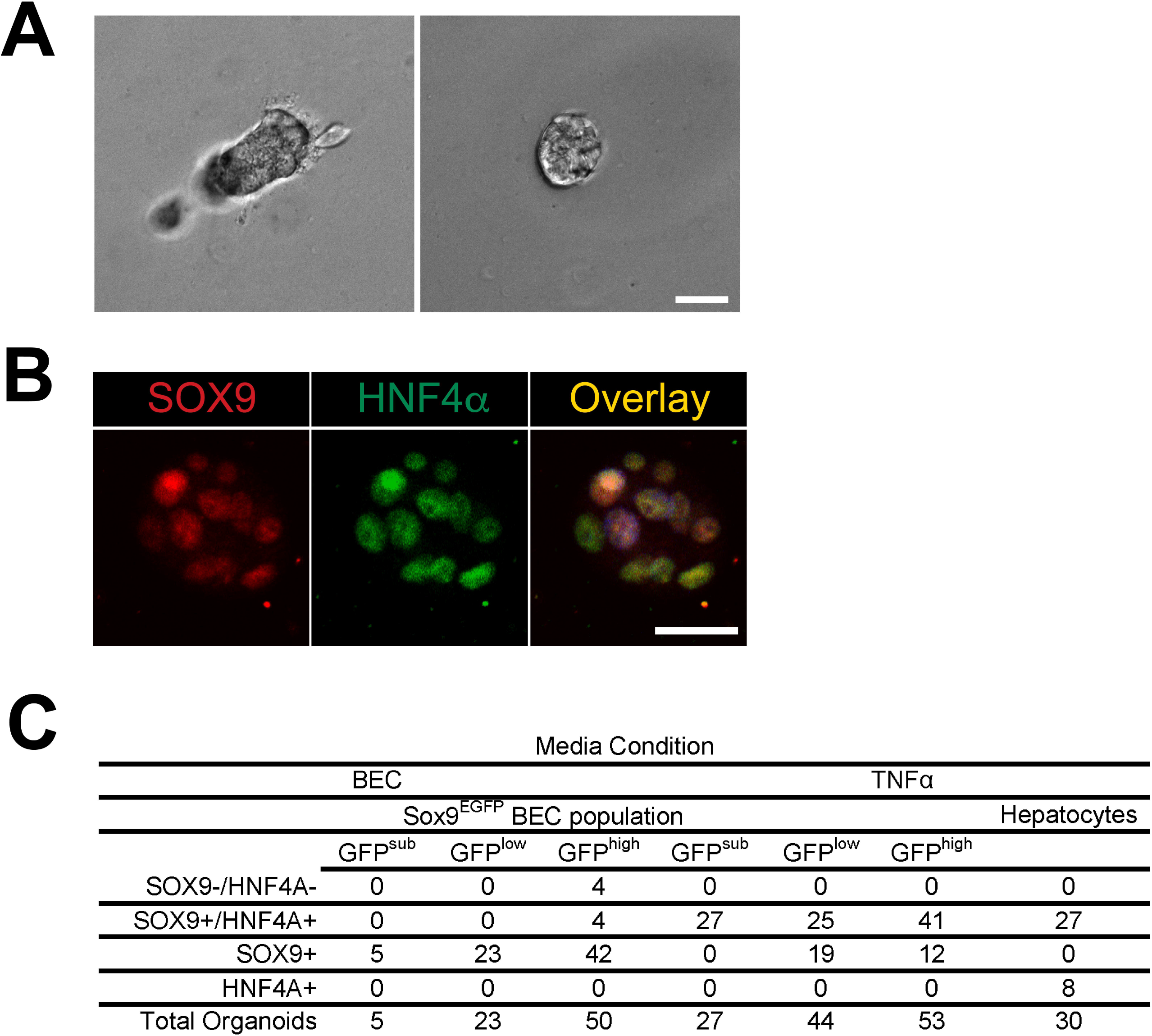
Expression of SOX9 and HNF4A in hepatocyte organoids. (A) Representative images of hepatocyte organoids following 7 days in culture (scale bar = 25μm). (B) A majority of hepatocyte organoids express both SOX9 and HNF4A *in vitro* (scale bar = 25μm). (C) Organoid quantification for SOX9/HNF4A co-localization (related to Fig. 6B).

**Supplemental Figure 5.**
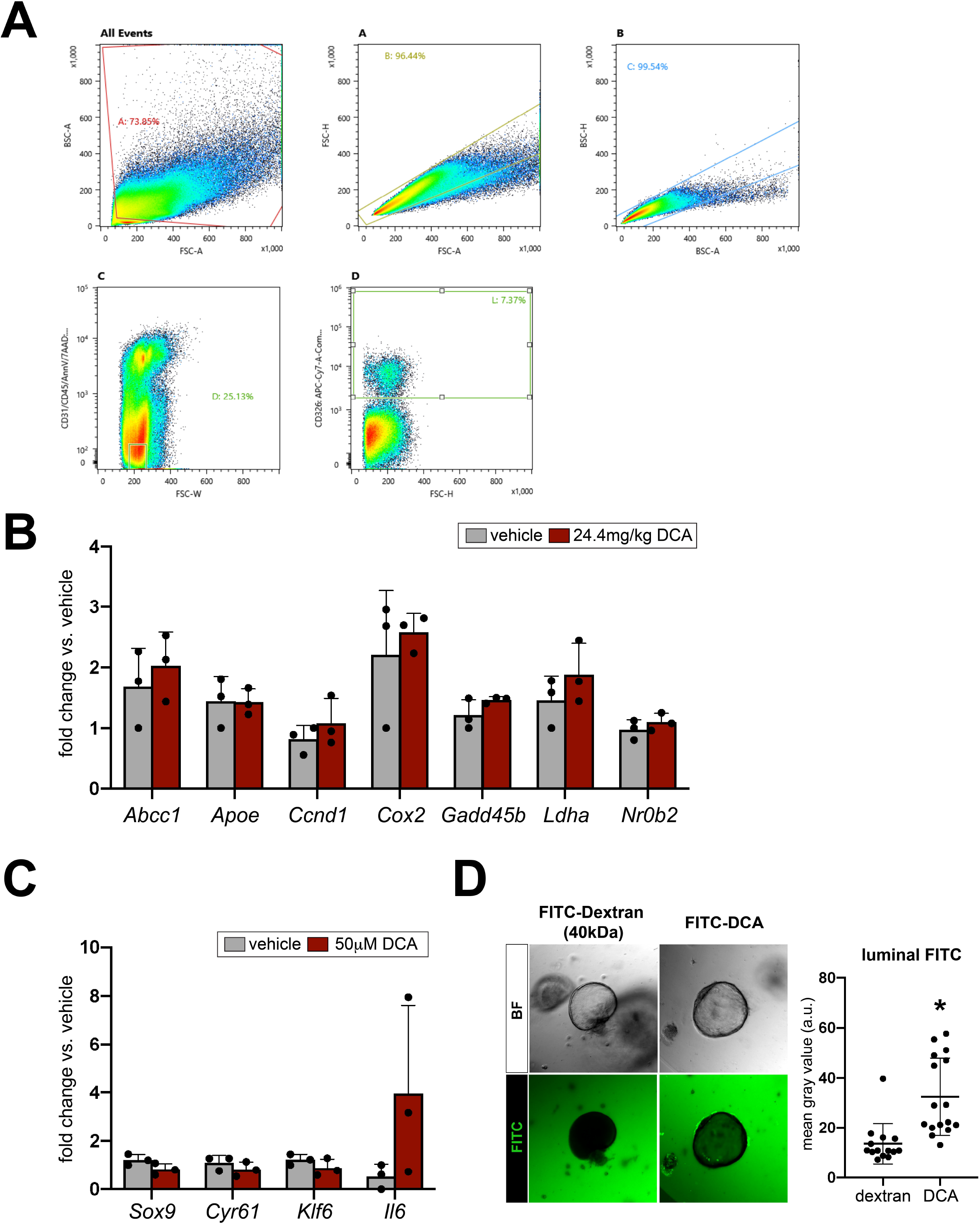
Biliary responses to deoxycholic acid. (A) FACS isolation strategy for EPCAM^+^ BECs from vehicle and DCA-treated mice. (B) Candidate DCA targets that were not differentially expressed in isolated BECs following DCA administration *in vivo*. (C) Biliary organoids treated with DCA for 24hr also failed to upregulate *Sox9*, *Cyr61*, and *Klf6* and demonstrated heterogeneous response in terms of *Il6* upregulation. (D) FITC-conjugated DCA was found in organoid lumens following 9hr of treatment, while FITC-dextran was excluded (n=16 FITC-dextran treated organoids, 14 FITC-DCA treated organoids; * indicates significance, p = 0.004, unpaired t-test).

**Supplemental Table 1. *Genes upregulated in a single Sox9^EGFP^ population by RNA-seq***

**Supplemental Table 2. *p values for gene set signature scores***

**Supplemental Table 3. *Antibodies***

**Supplemental Table 4. *Taqman probes***

## METHODS

### Mice

Sox9^EGFP^ mice were previously developed and characterized (Gong et al. 2003; Formeister et al. 2009). All Sox9^EGFP^ mice were heterozygous for the EGFP transgene and maintained on the C57Bl/6 background. All experiments were carried out on mice between 8-24 weeks of age. Male mice were used for gene expression studies, and male and female mice were used for immunofluorescence and organoid experiments. Mice were fed Tekland Global Soy Protein-Free Extruded Rodent Diet (Envigo, 2020SX) and received water *ad libitum*. For gene expression studies, mice were fasted overnight (12-14hr) prior to tissue harvest. For *in vivo* DCA administration, male C57Bl/6 mice were given 24.4mg/kg DCA in PBS (pH 7.4) or PBS only via a single intraperitoneal injection, as previously described (Paolini et al. 2002). DCA- and vehicle-treated mice were fasted 16hr and euthanized 24hr following treatment. Sox9^EGFP^ mice were phenotyped by observing EGFP expression in tail clippings by epifluorescent microscopy, as described in (Formeister et al. 2009). The Institutional Animal Care and Use Committee of the University of North Carolina at Chapel Hill reviewed and approved all animal use protocols.

### Immunofluorescence

Livers were fixed by intracardiac perfusion of 4% PFA in PBS, followed by incubation overnight in 4% PFA in PBS at 4°C. Tissue was transferred to 30% sucrose and incubated overnight at 4°C. Livers were dissected into individual lobes, embedded in Tissue-Tek Optimal Cutting Temperature media (Sakura), frozen, and cut into 10μm sections. Sections were stored at −80°C until use.

Immunofluorescence was carried out as previously described, and all steps were carried out at room temperature unless otherwise noted (Raab et al. 2020). Sections were permeabilized with 0.3% Triton X-100 in PBS for 20min, then blocked in 1X Animal-Free Blocking Solution (Cell Signaling Technologies, 15019) for 30min. Primary antibodies were applied in PBS for 2hr at room temperature or overnight at 4°C. Slides were rinsed in PBS, three times for 5min. Secondary antibodies were applied in PBS for 45min, and slides were rinsed in PBS, three times for 5min. Nuclei were stained with bisbenzimide (Sigma-Aldrich, B2883), diluted 1:1000 in PBS for 10min, prior to a final wash in PBS, three times for 5min. Slides were mounted to coverslips using Hydromount Mounting Media (Electron Microscopy Services, 17966). For semi-quantitative confocal analysis of EGFP, slides were mounted using ProLong Gold Mounting Media (Thermo Fisher, P36934). Antibodies used in this manuscript are listed in Supplemental Table 2. For co-labeling with WGA, tissue was incubated for 10min in 1mg/mL WGA-AlexaFluor647 (Thermo Fisher, W32466) in HBSS prior to permeabilization. Following WGA incubation, tissue was washed in HBSS, twice for 5min.

Images were acquired using an Olympus IX-81 widefield epifluorescent microscope or Zeiss LSM700 confocal microscope. Confocal images of tissue were acquired as 1μM optical sections; confocal images of organoids were acquired as 5 μM optical sections. Low-magnification image of Sox9^EGFP^ right lobe (Fig. 1A) was acquired by tile scan imaging of fresh tissue immediately following dissection. Confocal imaging was carried out in the Microscopy Services Laboratory at UNC Chapel Hill. Statistical analyses were carried out in Prism 8.4.2 (GraphPad).

### Intrahepatic bile duct isolation

To isolate single BECs from intrahepatic bile ducts, we dissociated liver tissue as previously described, with minor modifications (Broutier et al. 2016). Briefly, liver lobes were dissected without removing extrahepatic duct and gallbladder, to avoid contamination of intrahepatic biliary preps with extrahepatic biliary tissue. The tissue was rinsed in sterile DPBS (Gibco), minced (<0.5cm^2^ pieces) with a razor blade, and transferred to a 50mL conical tube. Liver tissue pieces were rinsed twice with Liver Wash Buffer [DMEM-H (Gibco), 1% FBS, 1% Glutamax (Gibco), 1% Pen/Strep (Gibco)] by resuspending and allowing tissue to sediment before removing supernatant. Following the second rinse, tissue was centrifuged at 600g at 4°C for 5 minutes to remove residual Liver Wash Buffer. The supernatant was discarded and liver tissue pieces were resuspended in 10 mL of Dissociation buffer [Liver Wash Buffer with 0.15mg/mL collagenase type XI (Sigma, C9407), 0.3U/mL Dispase (Corning, 354235), 200ug/mL DNase (Sigma DN25)], pre-warmed to 37°C. The tube was transferred to a rocking platform (60RPM) and incubated at 37°C for 90min with continuous shaking. The tube was retrieved every 30 minutes and vigorously pipetted 50 times using a P1000 micropipette set to 1mL, to aid in dissociation of bile ducts. After 90min of digest, dissociated tissue was rinsed twice with 10mL of Liver Wash Buffer followed by centrifugation at 200g at 4°C for 5 minutes. Supernatant was discarded after each wash by careful decanting, so as not to disturb the cell pellet. The pellet was resuspended in 5mL of Red Blood Cell Lysis Buffer (Biolegend, 420301) diluted in sterile Molecular Grade Water (Corning), as per manufacturer protocol, and incubated on ice for 5 minutes with repeated agitation. 10mL of DPBS (Gibco) was added to quench the lysis buffer and the tube was centrifuged at 600g at 4°C for 5min. Supernatant was discarded and ductal fragments were resuspended in 500uL Advanced DMEM (Gibco) for whole duct organoid culture, or used for single cell dissociation (see ***Single BEC dissociation and flow cytometry/FACS***, below).

### Single BEC dissociation and flow cytometry/FACS

Ductal fragments were resuspended in 1 mL of TrypLE (Gibco) with 100μg/mL Y-27632 (Selleck Chemicals, S1049) for single cell dissociation. The tube was incubated at 37°C for 12 minutes with vigorous pipetting every 2 minutes, to aid in dissociation of isolated ductal fragments. 10mL of Advanced DMEM (Gibco) was added to quench the TrypLE and stop the dissociation. Dissociated cells were passed through a 40μm cell strainer into a new 50mL conical tube and centrifuged at 600g at 4°C for 5min. The supernatant was discarded and cells were resuspended in 500μL of Sort Media [Advanced DMEM with 500μL B27 Supplement without Vitamin A (Thermo Fisher) - 250μL N2 Supplement (Thermo Fisher) – 250μL Glutamax (Gibco) – 250uL Hepes Buffer Solution (Gibco) – 250μL Pen/Strep (Gibco) – 0.1% Y-27632 100 μL/mL (Selleck Chemicals, S1049) – 0.1% DNase (Sigma DN25)].

Single BECs were stained with anti-CD31-APC (1:100) (Biolegend) and anti-CD45-APC (1:100) (Biolegend) and incubated, covered, on ice for 45min. Cells were rinsed with 3mL of Advanced DMEM (Gibco) and centrifuged at 600g at 4°C for 5 minutes. After decanting the supernatant, the cells were resuspended in 1.5 mL of Sort Media. 5μL of 7AAD (Biolegend, 420404) and 5μL of AnnexinV-APC (Biolegend, 640941) were added to resuspended cells to distinguish dead and dying cells, respectively. Cells were analyzed and collected using a Sony SH800 fluorescence-activated cell sorter. Gating schemes are shown in Supplemental Figures 2 and 5. For gene expression experiments, cells were collected directly into 500μL RNA Lysis Buffer (Ambion RNAqueous Micro Kit, AM1931); for organoid culture experiments, cells were collected directly into Sort Media.

### RNA isolation and RT-qPCR

FACS-isolated cells and organoids for gene expression studies were lysed in RNA Lysis Buffer and RNA was isolated using the RNAqueous Micro Kit (Ambion), following manufacturer instructions. cDNA was generated with the iScript cDNA Synthesis Kit (BioRad, 1708891) and diluted 1:10 in molecular grade H2O (Corning) for RT-qPCR. RT-qPCR was carried out using Taqman probes and SsoAdvanced Universal Probes Supermix (BioRad, 1725280). To detect nascent *Sox9* RNA, primers were designed spanning the junction of *Sox9* exon 2-intron 2 using Primer-BLAST (Ye et al. 2012). Primers were validated to have efficiency between 90-110% and produced a single PCR product that was validated by Sanger sequencing (Eurofins Genomics). RT-qPCR for nascent *Sox9* was carried out using SsoAdvanced Universal SYBR Green Supermix (BioRad, 1725270). Relative fold change of gene expression was calculated using the delta-delta C_T_ method, with *18S* as the internal reference gene (Pfaffl 2001). Statistical analyses were carried out in Prism 8.4.2 (GraphPad). Taqman assay IDs and *Sox9* primer sequences used in this manuscript are listed in Supplemental Table 3.

### RNA-seq and analysis

Libraries for RNA-seq were prepared as previously described (Raab et al. 2020). 12,000 cells per Sox9^EGFP^ population were collected directly into 500uL of RNA Lysis Buffer (Ambion RNAqueous Micro Kit) and RNA was isolated as described above (see ***RNA isolation and RT-qPCR***), with an adjustment to the protocol so that final volume of eluate is equivalent to 15mL. cDNA was prepared from 5mL of RNA, which was used to validate FACS isolation by RT-qPCR for *Egfp* (Life Technologies) using *18S* as the internal reference gene. 1mL of RNA from each sample was used to assess RNA integrity number by Bioanalyzer (Agilent) using the RNA Pico assay. RIN values were ≥ 7.0 for all RNA-seq samples. Libraries were prepared from 8mL of RNA using the SMARTer Stranded Total RNA-seq Pico kit v2 (Clontech), per manufacturer instructions. Libraries were pooled in equimolar ratios using concentrations determined by Qubit 3.0 High Sensitivity DNA assay (Life Technologies) and Bioanalyzer (Agilent) using the High Sensitivity DNA Analysis kit. Libraries were sequenced on a NextSeq500 (Illumina) with 75 bp single-end reads, v2 chemistry.

Transcript abundances were estimated using Salmon (1.1.0) indexed to Gencode mouse annotations v24 and collapsed to gene counts using tximport (1.14.2) (Soneson et al. 2015; Patro et al. 2017). Differential expression analysis was performed using DESeq2 (1.26.0) (Love et al. 2014). Heatmaps were generated using ComplexHeatmap (2.2.0) and the Z-scored normalized count values. Signature scores were derived using the singscore R package (1.6.0) (Foroutan et al. 2018); data from (Dong et al. 2007; Villanueva et al. 2012; Schaub et al. 2018). All code used in these analysis available at DOI: 10.5281/zenodo.3858321. RNA-seq data is deposited in GEO under series number: GSE151387.

### Primary hepatocyte isolation

Hepatocyte isolation was adapted from a previously published protocol (Kim et al. 2006). Briefly, mice weighing 20-35g were anesthetized with Nembutal (40-60 mg/kg, IP). Abdominal cavity was opened, and portal vein was cannulated (24G × ¾” Terumo™ Surflo™ ETFE I.V. catheter, Fisher Scientific, Hampton, NH). The catheter was connected to a perfusion system for cell isolation from mouse organs (Cat #73-3659, Harvard Apparatus, Holliston, MA). Liver was perfused with 50 ml of 37°C sterile buffer I solution [50 mM EGTA, 1M glucose, 1% pen/strep in calcium- and magnesium-free Hank’s Balanced Salt Solution (HBSS)] at a rate of 7 ml/min. At this time, inferior vena cava was cut. Subsequently, liver was digested with 40 mL of 37°C sterile buffer II solution [1M CaCl_2_, 1M glucose, 1% pen/strep, and 3600 U of Collagenase IV in calcium- and magnesium-free Hank’s Balanced Salt Solution (HBSS)] at a rate of 5 ml/min. Livers were surgically removed, and hepatocytes were released in cold isolation media (1X DMEM, 1% pen/strep) by removing the Glisson’s capsule using sterile tweezers. Hepatocytes were isolated by size exclusion using 100mm and 70mm filters, respectively. Cells were washed at 120 × *g* for 5 min at 4°C. Live/dead cell exclusion was performed by Percoll gradient (1:1 1X Percoll/isolation media) and centrifugation at 120g for 5 min at 4°C. Hepatocytes were resuspended in isolation media.

### Biliary and hepatocyte organoid culture

#### Whole duct culture and bile acid treatment

10μL aliquots from ductal preps (see ***Intrahepatic bile duct isolation***) were examined by light microscopy to qualitatively determine cellular density. 5-10μL of ductal prep was diluted in Advanced DMEM/F12 (Gibco) and Cultrex Type II Growth Factor Reduced extracellular matrix (R&D Biosystems 3533-010-02) was added to a final concentration of 66% Cultrex. Cultrex-duct suspensions were plated as 40μL droplets per well in pre-warmed 48-well plates and allowed to polymerize at 37°C for 20min. After polymerization, droplets were overlaid with 200μL Biliary Expansion Media [50% Advanced DMEM/F12 (Gibco), 40% WNT3A-conditioned media, 10% RSPO1-conditioned media, B27 Supplement without Vitamin A (Thermo Fisher), N2 Supplement (Thermo Fisher), Glutamax (Gibco), 10mM HEPES (Gibco), Penicillin/Streptomycin (Gibco), 50ng/mL recombinant murine EGF (Gibco, PMG8043), 100ng/mL recombinant human Noggin (Peprotech, 120-10C), 100ng/mL recombinant human FGF10 (Peprotech, 100-26), 10μM recombinant human Gastrin (Sigma-Aldrich G9145), 50ng/mL recombinant human HGF (Peprotech 100-39H), 10mM Nicotinamide, and 10μM Y-27632 (Selleck Chemicals)]. Media was replaced every other day. Starting on day 4, WNT3A-conditioned media was removed and replaced with Advanced DMEM/F12, and Noggin and Y-27632 were withdrawn from culture.

For bile acid and verteporfin organoid experiments, ductal organoids were passaged twice prior to treatment. Organoids were grown for 7 days prior to each passage, and passaged by removing media and adding 250uL TrypLE (Gibco). Cultrex droplets were mechanically dissociated in TrypLE by pipetting, then incubated at 37°C for 3min. TrypLE was quenched by adding 250uL Advanced DMEM/F12, and organoid fragments were pelleted at 6,000 × *g* for 5min at room temperature. Organoid fragments were re-plated at a 1:2 ratio as 40μL droplets (66% Culrex:34% Advanced DMEM/F12) in pre-warmed 48-well plates, as described above. For monolayer experiments, organoid fragments were passaged onto 48-well plates coated with 10% Cultrex at passage 2. Following each passage, organoids and monolayers were initially grown in Biliary Expansion Media with WNT3A, Noggin, and Y27632 for 4 days. WNT3A, Noggin, and Y27632 were removed from culture for 48hr prior to treatment. Samples were treated with 50μM deoxycholic acid (Sigma), 1μM verteporfin (Tocris, 5305), or an equivalent volume of DMSO for 24hr prior to lysis in 500uL of RNA Lysis Buffer (Ambion RNAqueous Micro Kit). To determine bile acid presence in organoid lumens, organoids were treated with 50μM FITC-conjugated deoxycholic acid (E. Mash, University of Arizona) or 40kDa FITC-conjugated dextran (Sigma, FD40S) (Jean-Louis et al. 2006). Organoid images were acquired as 1μM optical sections on a Zeiss LSM700 confocal microscope 9hr after treatment with deoxycholic acid or dextran.

### Single cell culture

FACS isolated BECs (see ***Single BEC dissociation and flow cytometry/FACS***) were resuspended in Cultrex Type II Growth Factor Reduced extracellular matrix (R&D Biosystems). For organoid survival and whole-mount immunofluorescence experiments, Cultrex-cell suspensions were plated as 40μL droplets per well in pre-warmed 48-well plates. For gene expression experiments, Cultrex-cell suspensions were plated as 10μL droplets per well to pre-warmed 96-well plates. All Cultrex droplets were allowed to polymerize at 37°C for 20min. After polymerization, BEC or TNFα media was overlaid: 200μL per well for 48-well plates or 100μL per well for 96-well plates. BECs were plated at density of 5,000 cells per 40μL Cultrex droplet per well (48-well plate) or 1,200 cells per 10uL Cultrex droplet per well (96-well plate). Primary hepatocytes were plated at a density of 10,000 cells per 40μL Cultrex droplet per well (48-well plate).

Organoids grown in BEC media were initially overlaid with Biliary Expansion Media. Media was replaced every other day. Starting on day 4, WNT3A-conditioned media was removed and replaced with Advanced DMEM/F12, and Noggin and Y-27632 were withdrawn from culture. Organoids grown in TNFα conditions were overlaid with TNFα Media [Advanced DMEM/F12 (Gibco), B27 Supplement without Vitamin A (Thermo Fisher), N2 Supplement (Thermo Fisher), Glutamax (Gibco), 10mM HEPES (Gibco), Penicillin/Streptomycin (Gibco), 25ng/mL recombinant murine EGF (Gibco), 50ng/mL recombinant human HGF (Peprotech), 10μM Y-27632 (Selleck Chemicals), 1μM A8301 (Tocris, 2939), 3μM CHIR99021 (Selleck Chemicals, S1263), and 100ng/mL recombinant murine TNFα (Peprotech, 315-01A). Media was replaced every other day.

#### Whole mount immunofluorescence

Organoids grown from single cells were fixed 7 days after plating, as follows. All steps were carried out at room temperature, unless noted. Overlay media was removed and 200μL of 4% paraformaldehyde (VWR 41678) in DPBS (pre-warmed at 37°C) was added to each well for 15min to fix the organoids. 4% paraformaldehyde was removed and the organoids were rinsed with 200μL PBS (Gibco) twice for 5min. The organoids were permeabilized with 200μL 0.5% Triton (Sigma-Aldrich T8787) in PBS for 20min, then rinsed with 200μL 100mM Glycine in PBS twice for 15min. Organoids were incubated with 250μL of Blocking Buffer [10% Normal Donkey Serum (Jackson Immunoresearch Laboratories 017-000-121) in Immunofluorescence Buffer (DPBS-0.1% Bovine Serum Albumin, 0.2% Triton, 0.05% Tween-20]] for 90min. Primary antibodies were applied in the 250μL of Blocking Buffer and incubated overnight at 4°C.

Primary antibodies were removed and organoids were washed with 250μL Immunofluorescence Buffer three times for 20min. Secondary antibodies were applied in 250μL of Blocking Buffer and incubated for 2hr. Secondary antibodies were removed and organoids were washed with 250μL Immunofluorescence Buffer three times for 20min. Nuclei were stained with a bisbenzimide (Sigma-Aldrich) diluted 1:1000 in Immunofluorescence Buffer for 30min. Organoids were washed with DPBS for 5min three times and immediately quantified. SOX9 and HNF4A expression was observed using an Olympus IX-81 inverted epifluorescent microscope.

## ACKNOWLEDGMENTS

The authors would like to thank Drs. Bailey Zwarcyz, Scott Magness, Susan Henning, Paul Dawson, Terry Magnuson, and members of the Magnuson Lab (UNC) for helpful discussion and feedback. FITC-DCA was the kind gift of Dr. Eugene A. Mash (University of Arizona).

## REFERENCES

Aloia L, McKie MA, Vernaz G, Cordero-Espinoza L, Aleksieva N, van den Ameele J, Antonica F, Font-Cunill B, Raven A, Aiese Cigliano R et al. 2019. Epigenetic remodelling licences adult cholangiocytes for organoid formation and liver regeneration. Nature cell biology 21: 1321–1333.

Alpini G, Roberts S, Kuntz SM, Ueno Y, Gubba S, Podila PV, LeSage G, LaRusso NF. 1996. Morphological, molecular, and functional heterogeneity of cholangiocytes from normal rat liver. Gastroenterology 110: 1636–1643.

Anakk S, Bhosale M, Schmidt VA, Johnson RL, Finegold MJ, Moore DD. 2013. Bile acids activate YAP to promote liver carcinogenesis. Cell Rep 5: 1060–1069.

Antoniou A, Raynaud P, Cordi S, Zong Y, Tronche F, Stanger BZ, Jacquemin P, Pierreux CE, Clotman F, Lemaigre FP. 2009. Intrahepatic bile ducts develop according to a new mode of tubulogenesis regulated by the transcription factor SOX9. Gastroenterology 136: 2325–2333.

Ayyaz A, Kumar S, Sangiorgi B, Ghoshal B, Gosio J, Ouladan S, Fink M, Barutcu S, Trcka D, Shen J et al. 2019. Single-cell transcriptomes of the regenerating intestine reveal a revival stem cell. Nature 569: 121–125.

Broutier L, Andersson-Rolf A, Hindley CJ, Boj SF, Clevers H, Koo BK, Huch M. 2016. Culture and establishment of self-renewing human and mouse adult liver and pancreas 3D organoids and their genetic manipulation. Nat Protoc 11: 1724–1743.

Dai J, Wang H, Dong Y, Zhang Y, Wang J. 2013. Bile acids affect the growth of human cholangiocarcinoma via NF-kB pathway. Cancer Invest 31: 111–120.

DeLeve LD, Wang X, Hu L, McCuskey MK, McCuskey RS. 2004. Rat liver sinusoidal endothelial cell phenotype is maintained by paracrine and autocrine regulation. American journal of physiology Gastrointestinal and liver physiology 287: G757–763.

Dong J, Feldmann G, Huang J, Wu S, Zhang N, Comerford SA, Gayyed MF, Anders RA, Maitra A, Pan D. 2007. Elucidation of a universal size-control mechanism in Drosophila and mammals. Cell 130: 1120–1133.

Font-Burgada J, Shalapour S, Ramaswamy S, Hsueh B, Rossell D, Umemura A, Taniguchi K, Nakagawa H, Valasek MA, Ye L et al. 2015. Hybrid Periportal Hepatocytes Regenerate the Injured Liver without Giving Rise to Cancer. Cell 162: 766–779.

Formeister EJ, Sionas AL, Lorance DK, Barkley CL, Lee GH, Magness ST. 2009. Distinct SOX9 levels differentially mark stem/progenitor populations and enteroendocrine cells of the small intestine epithelium. American journal of physiology Gastrointestinal and liver physiology 296: G1108–1118.

Foroutan M, Bhuva DD, Lyu R, Horan K, Cursons J, Davis MJ. 2018. Single sample scoring of molecular phenotypes. BMC Bioinformatics 19: 404.

Gong S, Zheng C, Doughty ML, Losos K, Didkovsky N, Schambra UB, Nowak NJ, Joyner A, Leblanc G, Hatten ME et al. 2003. A gene expression atlas of the central nervous system based on bacterial artificial chromosomes. Nature 425: 917–925.

Halpern KB, Shenhav R, Massalha H, Toth B, Egozi A, Massasa EE, Medgalia C, David E, Giladi A, Moor AE et al. 2018. Paired-cell sequencing enables spatial gene expression mapping of liver endothelial cells. Nat Biotechnol 36: 962–970.

Hu H, Gehart H, Artegiani B, C LO-I, Dekkers F, Basak O, van Es J, Chuva de Sousa Lopes SM, Begthel H, Korving J et al. 2018. Long-Term Expansion of Functional Mouse and Human Hepatocytes as 3D Organoids. Cell 175: 1591–1606 e1519.

Huch M, Dorrell C, Boj SF, van Es JH, Li VS, van de Wetering M, Sato T, Hamer K, Sasaki N, Finegold MJ et al. 2013. In vitro expansion of single Lgr5+ liver stem cells induced by Wnt-driven regeneration. Nature 494: 247–250.

Jean-Louis S, Akare S, Ali MA, Mash EA, Jr., Meuillet E, Martinez JD. 2006. Deoxycholic acid induces intracellular signaling through membrane perturbations. J Biol Chem 281: 14948–14960.

Kamimoto K, Kaneko K, Kok CY, Okada H, Miyajima A, Itoh T. 2016. Heterogeneity and stochastic growth regulation of biliary epithelial cells dictate dynamic epithelial tissue remodeling. Elife 5.

Kim ND, Moon JO, Slitt AL, Copple BL. 2006. Early growth response factor-1 is critical for cholestatic liver injury. Toxicol Sci 90: 586–595.

Liu-Chittenden Y, Huang B, Shim JS, Chen Q, Lee SJ, Anders RA, Liu JO, Pan D. 2012. Genetic and pharmacological disruption of the TEAD-YAP complex suppresses the oncogenic activity of YAP. Genes Dev 26: 1300–1305.

Love MI, Huber W, Anders S. 2014. Moderated estimation of fold change and dispersion for RNA-seq data with DESeq2. Genome Biol 15: 550.

Manco R, Clerbaux LA, Verhulst S, Bou Nader M, Sempoux C, Ambroise J, Bearzatto B, Gala JL, Horsmans Y, van Grunsven L et al. 2019. Reactive cholangiocytes differentiate into proliferative hepatocytes with efficient DNA repair in mice with chronic liver injury. J Hepatol 70: 1180–1191.

Maroni L, Haibo B, Ray D, Zhou T, Wan Y, Meng F, Marzioni M, Alpini G. 2015. Functional and structural features of cholangiocytes in health and disease. Cell Mol Gastroenterol Hepatol 1: 368–380.

Mederacke I, Hsu CC, Troeger JS, Huebener P, Mu X, Dapito DH, Pradere JP, Schwabe RF. 2013. Fate tracing reveals hepatic stellate cells as dominant contributors to liver fibrosis independent of its aetiology. Nat Commun 4: 2823.

Paolini M, Pozzetti L, Montagnani M, Potenza G, Sabatini L, Antelli A, Cantelli-Forti G, Roda A. 2002. Ursodeoxycholic acid (UDCA) prevents DCA effects on male mouse liver via up-regulation of CYP [correction of CXP] and preservation of BSEP activities. Hepatology 36: 305–314.

Patro R, Duggal G, Love MI, Irizarry RA, Kingsford C. 2017. Salmon provides fast and bias-aware quantification of transcript expression. Nat Methods 14: 417–419.

Peng WC, Logan CY, Fish M, Anbarchian T, Aguisanda F, Alvarez-Varela A, Wu P, Jin Y, Zhu J, Li B et al. 2018. Inflammatory Cytokine TNFalpha Promotes the Long-Term Expansion of Primary Hepatocytes in 3D Culture. Cell 175: 1607–1619 e1615.

Pepe-Mooney BJ, Dill MT, Alemany A, Ordovas-Montanes J, Matsushita Y, Rao A, Sen A, Miyazaki M, Anakk S, Dawson PA et al. 2019. Single-Cell Analysis of the Liver Epithelium Reveals Dynamic Heterogeneity and an Essential Role for YAP in Homeostasis and Regeneration. Cell Stem Cell.

Pfaffl MW. 2001. A new mathematical model for relative quantification in real-time RT-PCR. Nucleic Acids Res 29: e45.

Planas-Paz L, Sun T, Pikiolek M, Cochran NR, Bergling S, Orsini V, Yang Z, Sigoillot F, Jetzer J, Syed M et al. 2019. YAP, but Not RSPO-LGR4/5, Signaling in Biliary Epithelial Cells Promotes a Ductular Reaction in Response to Liver Injury. Cell Stem Cell 25: 39–53 e10.

Poling HM, Mohanty SK, Tiao GM, Huppert SS. 2014. A comprehensive analysis of aquaporin and secretory related gene expression in neonate and adult cholangiocytes. Gene Expr Patterns 15: 96–103.

Raab JR, Tulasi DY, Wager KE, Morowitz JM, Magness ST, Gracz AD. 2020. Quantitative classification of chromatin dynamics reveals regulators of intestinal stem cell differentiation. Development 147.

Ramalingam S, Daughtridge GW, Johnston MJ, Gracz AD, Magness ST. 2012. Distinct levels of Sox9 expression mark colon epithelial stem cells that form colonoids in culture. American journal of physiology Gastrointestinal and liver physiology 302: G10–20.

Raven A, Lu WY, Man TY, Ferreira-Gonzalez S, O’Duibhir E, Dwyer BJ, Thomson JP, Meehan RR, Bogorad R, Koteliansky V et al. 2017. Cholangiocytes act as facultative liver stem cells during impaired hepatocyte regeneration. Nature 547: 350–354.

Rezanejad H, Ouziel-Yahalom L, Keyzer CA, Sullivan BA, Hollister-Lock J, Li WC, Guo L, Deng S, Lei J, Markmann J et al. 2018. Heterogeneity of SOX9 and HNF1beta in Pancreatic Ducts Is Dynamic. Stem Cell Reports 10: 725–738.

Rimland CA, Tilson SG, Morell CM, Tomaz RA, Lu WY, Adams SE, Georgakopoulos N, Otaizo-Carrasquero F, Myers TG, Ferdinand JR et al. 2020. Regional differences in human biliary tissues and corresponding in vitro derived organoids. Hepatology.

Sampaziotis F, de Brito MC, Madrigal P, Bertero A, Saeb-Parsy K, Soares FAC, Schrumpf E, Melum E, Karlsen TH, Bradley JA et al. 2015. Cholangiocytes derived from human induced pluripotent stem cells for disease modeling and drug validation. Nat Biotechnol 33: 845–852.

Schaub JR, Huppert KA, Kurial SNT, Hsu BY, Cast AE, Donnelly B, Karns RA, Chen F, Rezvani M, Luu HY et al. 2018. De novo formation of the biliary system by TGFbeta-mediated hepatocyte transdifferentiation. Nature 557: 247–251.

Soneson C, Love MI, Robinson MD. 2015. Differential analyses for RNA-seq: transcript-level estimates improve gene-level inferences. F1000Res 4: 1521.

Song S, Ajani JA, Honjo S, Maru DM, Chen Q, Scott AW, Heallen TR, Xiao L, Hofstetter WL, Weston B et al. 2014. Hippo coactivator YAP1 upregulates SOX9 and endows esophageal cancer cells with stem-like properties. Cancer Res 74: 4170–4182.

Thomas AM, Hart SN, Kong B, Fang J, Zhong XB, Guo GL. 2010. Genome-wide tissue-specific farnesoid X receptor binding in mouse liver and intestine. Hepatology 51: 1410–1419.

Troeger JS, Mederacke I, Gwak GY, Dapito DH, Mu X, Hsu CC, Pradere JP, Friedman RA, Schwabe RF. 2012. Deactivation of hepatic stellate cells during liver fibrosis resolution in mice. Gastroenterology 143: 1073–1083 e1022.

Villanueva A, Alsinet C, Yanger K, Hoshida Y, Zong Y, Toffanin S, Rodriguez-Carunchio L, Sole M, Thung S, Stanger BZ et al. 2012. Notch signaling is activated in human hepatocellular carcinoma and induces tumor formation in mice. Gastroenterology 143: 1660–1669 e1667.

Ye J, Coulouris G, Zaretskaya I, Cutcutache I, Rozen S, Madden TL. 2012. Primer-BLAST: a tool to design target-specific primers for polymerase chain reaction. BMC Bioinformatics 13: 134.

Yi J, Lu L, Yanger K, Wang W, Sohn BH, Stanger BZ, Zhang M, Martin JF, Ajani JA, Chen J et al. 2016. Large tumor suppressor homologs 1 and 2 regulate mouse liver progenitor cell proliferation and maturation through antagonism of the coactivators YAP and TAZ. Hepatology 64: 1757–1772.

Yimlamai D, Christodoulou C, Galli GG, Yanger K, Pepe-Mooney B, Gurung B, Shrestha K, Cahan P, Stanger BZ, Camargo FD. 2014. Hippo pathway activity influences liver cell fate. Cell 157: 1324–1338.

Zong Y, Stanger BZ. 2012. Molecular mechanisms of liver and bile duct development. Wiley Interdiscip Rev Dev Biol 1: 643–655.

